# The ParA-like ATPase PldP influences the TatA dynamics in *Corynebacterium glutamicum*

**DOI:** 10.1101/2025.07.02.662783

**Authors:** Ekaterina Karnaukhova, Dominik Alwardt, Kati Böhm, Manuela Weiß, Fabian M. Meyer, Giacomo Giacomelli, Denise Mehner-Breitfeld, Thomas Brüser, Marc Bramkamp

## Abstract

In bacterial cells, precise localization of protein complexes is achieved by unique positioning systems. One of the examples of such positioning systems is the ParAB*-parS* which is responsible for plasmid and chromosome segregation. In *Corynebacterium glutamicum*, a *parAB* deletion results in cell division and growth defects, while deletion of an orphan ParA-like protein *pldP* results only in a moderate cell division phenotype. Having confirmed a basal ATPase activity of PldP, we aimed to explore if the Δ*pldP*-related phenotype could be a consequence of the mislocalized secreted proteins, as the loss of extracellular proteins involved in cell wall metabolism results in a similar phenotype characterized by disrupted separation of daughter cells. Putative peptidoglycan hydrolase Rv2525c from *Mycobacterium tuberculosis* and Rv2525c-like glycoside hydrolase-like domain-containing protein Cg0955 from *C. glutamicum* were previously shown to be transported outside of the cell by twin-arginine protein translocation machinery (Tat). Here, we found that although the deletion of *pldP* did not lead to the altered secretion of the putative hydrolase Cg0955 by the Tat system, it resulted in the reduction of the Tat dynamics. Our findings highlight the interplay between the ParA-like ATPase PldP and the Tat translocon and contribute to the studies of ParA-like proteins being essential in positioning various cargos in the bacterial cells.

**Importance:** Precise spatio-temporal localization of protein complexes within a bacterial cell is essential for the survival and proliferation of bacteria. ParA-like ATPases play a crucial role in protein positioning, as well as chromosome and plasmid segregation. Here, we characterize a novel ParA-like ATPase PldP in *Corynebacterium glutamicum*, a model organism for the cell biology of Mycobacteriales and a biotechnological workhorse. Deletion of *pldP* results in the cell division phenotypes and impacts the intracellular dynamics of TatA, a component of the twin-arginine protein transport. We suggest that the mislocalization of the Tat-secreted putative peptidoglycan hydrolase caused by the indirect influence of *pldP* deletion might account for the observed cell separation defect.

## Introduction

Life requires a precise spatial and temporal organization of biochemical processes. This fundamental principle becomes particularly obvious in processes such as cell elongation and maintenance of cell shape (1–4), chemotaxis (5–8), and differentiation (9), but also in membrane protein insertion and protein transport. All these processes rely on unique systems that ensure their timely and spatially correct functioning. However, bacterial cells often employ positioning systems that are based on a ParA-like ATPase (10, 11). ParA belongs to the group of the deviant Walker A motif ATPases (12, 13). The Walker A motif is responsible for ATP-binding. Unlike the classical motif, the deviant Walker A motif is characterized by a signature lysine residue at the beginning of the conserved sequence (XKGGXXK). The archetype of these dedicated partitioning proteins is the DNA-segregation ATPase ParA. ParA (derived from *par*titioning) proteins were first identified as plasmid segregation systems (13, 14). However, many chromosomes also encode ParA-based segregation systems that are mainly functioning in origin segregation. DNA partitioning systems comprise three components. The segregated DNA contains a recognition sequence termed *parS* sites (15, 16). These inverted repeats are bound by a ParB protein that acts as an adapter. During DNA segregation, ParB binds to *parS* to form a nucleoprotein complex, and this complex in turn stimulates ParA nucleotide hydrolysis (17, 18). The movement of ParB-*parS* is determined by the hydrolytic activity of the ATPase ParA. In the dimeric state, ParA-ATP is bound to DNA in a non-specific manner. By interacting with ParB-*parS*, ParA hydrolyzes ATP and dissociates from DNA, thus creating a gradient of ParA-ATP, in which the ParB-*parS* complex moves based on a diffusion ratchet mechanism (17, 19).

ParA-like ATPases not only function in the segregation of chromosomes and plasmids, but also in the positioning of carboxysomes (McdA in cyanobacteria (20)), and chemotaxis protein cluster (PpfA in *Rhodobacter sphaeroides* (21)). ParA, McdA, and PpfA bind to the nucleoid in a non-specific manner and use it as a matrix for movement. In contrast, in many rod-shaped bacteria, the ParA-like ATPase MinD binds to the cell membrane as a matrix (10, 22). MinD functions together with MinC to restrict Z-ring formation to the midcell (22). These findings highlight the diverse functions that ParA-like ATPases may have in partitioning.

The Gram-positive model bacterium *Corynebacterium glutamicum* contains a conventional *parAB* operon (*cg3427*, *cg3426* (23)). Deletions of *parA* and *parB* cause a decrease in cell growth and result in a high number of anucleate cells due to improper septum positioning (24). Additionally, we identified an orphan ParA-like protein, PldP (Cg1610) in *C. glutamicum* (23). Deletion of *pldP* leads to moderate cell length phenotypes, indicating that cell division or daughter cell separation does not work with normal precision in the mutant cells. In contrast to a *parA* deletion, the *pldP* deletion causes only a moderate increase in the number of anucleate cells (24). Until now, the function of PldP remained enigmatic (23). Corynebacteria are members of the actinobacteria phylum, and members of this phylum often share conserved cell biological features. Detailed literature research revealed that in *Streptomyces coelicolor*, the dynamic positioning of the twin-arginine protein translocation machinery (Tat) to the cell poles suggested an active localization (25). During aerial hypha formation, Tat proteins relocalize to the prespore compartments, which implies that there might be a system segregating the Tat components. Although the underlying mechanism could not be elucidated, the authors speculated about a ParA-like positioning system (25). The Tat pathway transports the folded proteins across the cell membrane (26, 27). In *C. glutamicum*, the components of the Tat pathway include TatA (Cg1685), TatB (Cg1273), TatC (Cg1684), and TatE (Cg3381) which is 71 % identical to TatA (28, 29). The *tatA* and *tatC* genes are encoded by the same operon, whereas *tatB* and *tatE* are monocistronic at distant chromosomal loci. Despite the differences in the genetic organization in *E. coli* and other Pseudomonadota, in which *tatB* and *tatC* are generally coupled in operons, all TatABC components of *C. glutamicum* are similarly essential for the formation of active Tat systems (30). Interestingly, in *E. coli,* the murein hydrolases AmiA and AmiC that are required for proper cell division are exported by the Twin-arginine transport (31), and, notably, also in *E. coli*, the Tat systems have been shown to be selectively positioned at the cell poles (32, 33). Therefore, we hypothesized that the cell division phenotype observed in a *pldP* deletion in *C*. *glutamicum* might well be attributed to a mislocalization of the Tat translocon.

In this study, we therefore examined whether the partitioning of Tat components in *C. glutamicum* depends on PldP. We hypothesized that the aberrant cell division of Δ*pldP* cells might be a consequence of improper Tat pathway function due to impaired positioning of the Tat components in the absence of PldP. Here, we show that PldP is an active ATPase and that its localization changes within the cell cycle. As evidenced by single-particle tracking, TatA proteins are dynamic, and their overall mobility decreases upon *pldP* deletion. However, we found that Tat-mediated protein secretion was not significantly affected in Δ*pldP* cells, indicating that the specific localization of the Tat system is not a general prerequisite for its function. In agreement with this, a direct interaction of PldP and Tat proteins could not be observed, and thus, the effect of PldP on Tat dynamics might be indirect. In summary, our findings confirm that PldP is an active ATPase that influences TatA dynamics. Although the overall transport of secreted test substrates is not affected by *pldP* deletion, the observed decrease in TatA dynamics hints at changes in membrane organization of the Tat secretion machinery that might indirectly cause the morphology phenotypes of a *pldP* mutant strain of *C. glutamicum*.

## Materials and Methods

### Growth medium and conditions

*E. coli* cells were grown at 37 °C in lysogeny broth (LB) containing if needed, kanamycin [25 µg/mL], or carbenicillin [100 µg/mL] and chloramphenicol [34 µg/mL]. *C. glutamicum* RES167 cells were grown at 30 °C in brain heart infusion (BHI) supplemented with kanamycin and IPTG [10, 50 or 100 µM], where appropriate. For growth analysis, *C. glutamicum* RES167 cells were grown overnight, and then the cultures were diluted to an optical density (OD_600_) of 0.1, and their growth was measured.

### Oligonucleotides, plasmids and bacterial strains

Oligonucleotides, plasmids and bacterial strains used in this study are listed in Tables S1-S3.

### Strain construction

All plasmid constructs were verified by sequencing (Eurofins Genomics). For pET16b and pK19*mobsacB* sequencing, oligonucleotides DA005, DA006, and GAA57, GAA58 were used, respectively. *Escherichia coli* NEB® 5-alpha (New England Biolabs) was used for cloning and plasmid propagation. For experiments, strains *E. coli* BL21(DE3)/pLysS (Promega) and *C. glutamicum* RES167, a restriction system-deficient derivative of the reference strain ATCC13032 (34, 35), were transformed with the indicated plasmids.

For inducible heterologous expression, the *pldP* gene (*cg1610*) was amplified from *C. glutamicum* gDNA via PCR and ligated into pET16b to generate plasmid pET16b-*pldP* (pAS001). Genes encoding a PldP with expected defects in ATP-binding (PldP^K50A^) or ATP-hydrolysis (PldP^D74A^) were obtained by performing site-directed mutagenesis (Q5 Site-Directed Mutagenesis Kit, NEB) in *pldP* using oligonucleotides DA001-DA004. It resulted in the generation of plasmids pET16b-*pldP^K50A^* (pDA001) and pET16b-*pldP^D74A^*(pDA002). Plasmids pAS001, pDA001 and pDA002 were used to transform *E. coli* BL21(DE3)/pLysS. Finally, strains EAD18-EAD20 were used to induce heterologous expression of *pldP*, *pldP^K50A^*, and *pldP^D74A^*, respectively. For inducible heterologous expression of *parB* (*cg3426*), the *parB* gene was amplified from *C. glutamicum* gDNA via PCR using oligonucleotides KB001 and KB002 and subsequently ligated into pET16b to generate plasmid pET16b-*parB* (pKB001). Plasmid pKB001 was transformed into *E. coli* BL21(DE3)/pLysS to generate strain EPH04. Positive clones were selected by carbenicillin and chloramphenicol, and confirmed by colony PCR.

For allelic exchange in *C. glutamicum* RES167, mobilization plasmid pK19*mobsacB* was used. In order to generate strains expressing *pldP::pldP-eYFP*, 500 bp of the *pldP* gene sequence, 500 bp of the downstream region, as well as the *eYFP* sequence, were amplified via PCR using oligonucleotides KB003-KB007. After restriction digestion, the fragments were ligated into pK19*mobsacB*, resulting in the plasmid pK19*mobsacB-pldP-eYFP* (pKB002). Plasmid pK19*mobsacB*-Δ*pldP* (pCD121) for *pldP* knock-out was generated as previously described (23). For strains expressing *tatA::tatA-halotag*, 500 bp of the *tatA* (*cg1685*) and its upstream region, 500 bp of the downstream region, and the *halotag* sequence were amplified via PCR using oligonucleotides EK43-EK48. After respective restriction digestion, the fragments were ligated into pK19*mobsacB*, generating plasmid pK19*mobsacB-tatA-halotag* (pEK01). This cloning strategy resulted in the construction of a strain synthesizing a TatA-HaloTag fusion as the only copy. Plasmid pK19*mobsacB*-*tatA-mCherry* (pDA003) was obtained by PCR amplification of the respective fragments using oligonucleotides DA007-DA014 and by ligating them via Gibson Assembly (Gibson Assembly Master Mix, NEB) into pK19*mobsacB*.

Before transformation, *C. glutamicum* RES167 cells were made competent according to a standard protocol (36). Purified mobilization plasmids pKB002, pKB003, pCD121, pEK01 and pDA003 were transformed into *C. glutamicum* RES167 cells via electroporation (37). The allelic exchange procedure was followed as previously described (38, 39). In short, the integration of the pK19*mobsacB* plasmid into the genome by a first crossover event was selected by kanamycin. Excision of the pK19*mobsacB* backbone was selected by sucrose [10 %]. The successful allelic exchange was confirmed by colony PCR. In summary, strains RES167 *pldP::pldP-eYFP* (CBK075), RES167 Δ*pldP* (CDC002), RES167 *tatA::tatA-halotag* (CEK01), and RES167 *tatA::tatA-mCherry* (CAD05) were constructed. Strain RES167 Δ*pldP tatA::tatA-halotag* (CEK02) was obtained by transforming plasmid pCD121 into strain CEK01. Strain RES167 Δ*pldP tatA::tatA-mCherry* (CAD06) was obtained by transforming plasmid pDA003 into strain CDC002.

Production of PldP-eYFP in CBK075 or HaloTag in strains CEK01 and CEK02 was confirmed via SDS-PAGE and following in-gel fluorescence using a ChemiDoc^TM^ MP Imaging System and the Image Lab Software (Bio-Rad). The presence of mCherry protein fusions in strains CAD05 and CAD06 was validated by Western blotting using anti-mCherry polyclonal rabbit antibodies (BioVision Inc.) in a 1:2,000 dilution. A secondary anti-rabbit polyclonal goat antibody conjugated to alkaline phosphatase (Sigma-Aldrich) was used in a 1:10,000 dilution. Secondary antibodies were detected with chromogenic alkaline phosphatase substrates 5-bromo-4-chloro-3-indolyl phosphate/nitro blue tetrazolium (Pierce^TM^ ECL Western Blotting Substrate, ThermoFisher Scientific).

### Genetic construction of Tat substrates for transport measurements

For analysis of Tat transport, the constitutive expression vectors pCLTON1-*P_sod_-cg0955-H6* (pDMB-RR-0955-H6), pCLTON1-*P_sod_-KK-cg0955-H6* (pDMB-KK-0955-H6), pCLTON1-*P_sod_-phoD-H6* (pDMB-RR-PhoD-H6) and pCLTON1-*P_sod_-KK-phoD-H6* (pDMB-KK-PhoD-H6) were constructed. For that, the P_sod_ promoter and the Cg0955-encoding gene of *C. glutamicum* (with stop codon exchanged by BamHI site) were first cloned into pUC19 (40), using the primers DMB001 (AatII) / DMB002 (AvrII) and DMB003 (AvrII) / DMB004 (BamHI) and the respective restriction sites. The XmnI-BamHI region of that vector was then cloned into pCLTON1 (41), thereby exchanging its *tetR*/P_tet_ region with the constitutive P_sod_/*phoD* expression system. The His6-tag and the *amiE* terminator were amplified with the pSG1154 template (42) and cloned in frame to the C-terminal end of *cg0955*, using BamHI/SacI restriction sites, resulting in pDMB-RR-0955-H6. For the construction of pDMB-RR-PhoD-H6, a backbone AvrII restriction site had to be removed from the vector by QuikChange mutagenesis (Agilent), using the primer DMB005 together with a primer of the corresponding complementary sequence, and the Cg0955-encoding gene was exchanged by *phoD* using the primers DMB006 (AvrII) / DMB007 (BamHI) for *phoD* amplification with genomic DNA as template, and the AvrII/BamHI restriction sites. Twin-arginines were exchanged by twin-lysines using QuikChange with the primers DMB008 (for the substrate Cg0955) or DMB009 (for the substrate PhoD), together with their respective complementary primers, resulting in the plasmids pDMB-KK-0955-H6 and pDMB-KK-PhoD-H6, respectively. *C. glutamicum* RES167 and Δ*pldP* cells were transformed with the plasmids, resulting in the generation of CDMB01-08 strains.

For gene overexpression in *C. glutamicum* RES167, the IPTG-inducible plasmid pEKEx2 (43) was used. For the construction of pEKEx2-*tatSP-mNeonGreen* (pEK02), the Tat signal peptide from the gene *cg0955*, and *mNeonGreen* were amplified via PCR using oligonucleotides EK39, EK40, and EK41, EK42, respectively. After restriction digestion, the fragments were ligated into pEKEx2, generating plasmid pEKEx2-*tatSP-mNeonGreen* (pEK02). Plasmid pEK02 was transformed into *C. glutamicum* RES167 and CDC002, generating strains RES167 pEKEx2-*tatSP-mNeonGreen* (CEK03) and RES167 Δ*pldP* pEKEx2-*tatSP-mNeonGreen* (CEK04), respectively. Kanamycin-resistant positive clones were selected and confirmed by colony PCR and mNeonGreen fluorescence.

### Protein purification

For heterologous expression of *pldP*, *pldP^K50A^*, *pldP^D74A^*and *parB*, strains EAD18-EAD20 and EPH04 were grown in 1 L of LB in the presence of chloramphenicol and carbenicillin. After OD_600_ reached 0.8, heterologous expression was induced with IPTG [0.4 mM], and the cultures were incubated for 4 h at 37 °C. Cells were harvested at 4,000 x g for 15 min at 4°C, frozen and stored at −80 °C until needed. Consequently, cell pellets were resuspended in Wash Buffer [50 mM Tris-HCl pH 7.5, 250 mM NaCl] supplemented with cOmplete^TM^ EDTA-free protease inhibitor cocktail and DNase I (Roche). Cells were lysed via French pressure cell press (SLM-AMINCO Spectronic Instruments). The suspension was passed through the press five times at 1,280 psi (20,000 psi inner cell pressure) and then centrifuged at 17,700 x g for 20 min at 4 °C. The proteins of interest were purified via Ni-nitriloacetic acid (Ni-NTA) affinity chromatography using ÄKTApurifier (Cytiva) and 1 mL Ni-NTA columns (Macherey-Nagel). The supernatant was passed through the Ni-NTA column which was subsequently washed in a step gradient of concentrations of Buffer A [50 mM Tris-HCl pH 7.5, 250 mM NaCl, 10 % glycerol, 5 mM MgCl_2_, 1 mM dithiothreitol, 10 mM imidazole] and Buffer B [same as Buffer A, but with 500 mM imidazole]. The step gradient was performed in four steps: 0, 5, 25, and 100 % of Buffer B. Peak fractions were pooled together and analyzed via SDS-PAGE, and the protein concentrations were measured via Bradford protein assay.

### ATPase activity assay

ATPase activities of PldP, PldP^K50A^ and PldP^D74A^ were tested via EnzChek^TM^ phosphate assay (Invitrogen), as described in the manual. Reaction volumes were reduced to 100 µL. The following concentrations were used for each reaction depending on the conditions: 5 µM PldP; 5 mM MgCl_2_; 0, 0.05, 0.1, 0.2, 0.5, 1, 2, and 5 mM ATP; 0, 0.1, 0.25, 0.5 and 0.75 mg/mL salmon sperm DNA (ThermoFisher Scientific); 0, 0.1 and 0.2 mg/mL liposomes. Liposomes were prepared from the *E. coli* total lipid extract (Avanti), as described previously (44). Lipids were dried, then resuspended and rehydrated in the gel filtration buffer C [50 mM HEPES-KOH, 150 mM KCl, 10 % glycerol, 0.1 mM EDTA]. Finally, lipids were extruded 40 times through 100 nm polycarbonate membranes (NucleporeTM Track-Etched Membrane 0.1 µm, Whatman). Other necessary components for the phosphate assay were added accordingly to the manual. Before adding ATP, the reactions were incubated for 10 min at 30 °C. Phosphate release was measured for each reaction at 30 °C for 3 h in the microplate reader at 360 nm (Tecan). After measurements, absorbance values for reactions with PldP were normalized to the background (controls without ATP, without DNA or liposomes) and to the ATP and DNA hydrolysis controls. These values were then compared to the phosphate standard curve in order to determine the phosphate number in nanomoles released by the ATPase activity of PldP. Data were analyzed with RStudio (Posit, PBC) using packages readxl (45), tidyr (46), drc (47), ggplot2 (48), and ggthemes (49).

ATPase activity of PldP was measured in the presence of ParB via Malachite Green Phosphate Assay Kit (Sigma-Aldrich) according to the manual. For this assay, 5 µM PldP was incubated with 3.5 mM ATP and increasing ParB concentrations [0, 5, 15, 30, 40, 50 µM]. ATPase assay was performed for 30 min at 30 °C, and the reaction was stopped with the Malachite Green reagent. The photometric detection was carried out using an Ultraspec 3000 photometer at 620 nm (Pharmacia Biotech). The data analysis was performed as described above.

### Fluorescence microscopy

Overnight cultures of *C. glutamicum* RES167 were diluted to an OD_600_ of 0.5 and grown until an OD_600_ of ∼1.8 in BHI supplemented with kanamycin and IPTG if necessary. 1 mL of cells was harvested and resuspended in 1X PBS [137 mM NaCl, 2.7 mM KCl, 8 mM Na_2_HPO_4_ x 2 H_2_O, 1.5 mM KH_2_PO_4_, pH 7.4]. Cells were mounted on agarose-coated slides [1 % agarose], and images were acquired via a Zeiss Axio Imager Microscope equipped with an EC Plan-Neofluar 100X/1.3 Oil Ph3 objective and an Axiocam camera (Zeiss). In the case of strains CEK01 and CEK02, before imaging, harvested cells were resuspended in BHI, and the HaloTag ligand TMR [50 nM] (Promega) was added. Cells were incubated at 30 °C for 1 h. In order to stain nucleoids, Hoechst 33342 (Thermo Scientific) was added to the samples for 5 min. After staining, cells were harvested, washed three times in 1X PBS, and imaged. In the case of strains CEK03 and CEK04, cell membranes were additionally stained with FM^TM^4-64 (Invitrogen) for 5 min, and resuspended in 1X PBS. TMR and FM^TM^4-64 molecules were excited at 558 nm, Hoechst 33342 – 359 nm, mNeonGreen – 470 nm, eYFP – 513 nm. TMR and FM^TM^4-64 emissions were measured at 583 nm, Hoechst 33342 – 461 nm, mNeonGreen – 509 nm, eYFP – 527 nm.

Obtained images were analyzed via Fiji (50). To detect differences in mNeonGreen fluorescence between strains CEK03 and CEK04, a set of custom Fiji macros was applied, as previously described (51). Firstly, phase contrast and fluorescence channels were aligned via a custom macro “Translate”. Secondly, three parameters were measured for each cell: total mNeonGreen fluorescence intensity of a cell and the fluorescence intensities at cell poles and septum (radius of measurement was 13 pixels). Fluorescence was represented as integrated density. Subsequent statistical analysis of the obtained values was performed via RStudio and GraphPad Prism (Dotmatics).

### Fluorescence recovery after photobleaching (FRAP)

Overnight cultures of CAD05 and CAD06 strains were diluted to an OD_600_ of 0.5 and grown at 30 °C, 200 rpm until an OD_600_ of ∼2 was reached. 1 ml of cells was harvested and resuspended in 1X PBS. Cells were mounted on agarose-coated slides and imaged via an Axio Observer.Z1 / 7 equipped with a Plan-Apochromat 40x/1.4 Oil DIC (UV) VIS-IR M27 objective (Zeiss). mCherry was excited at 587 nm, and emission was recorded at 610 nm. Two imaging devices were used: an electronically switchable illumination and detection module (ESID) and an Airyscan imaging device with a sensitivity set to 850 V. The power of the laser was set to 0.2% for imaging and to 100 % for the photobleaching of mCherry. For photobleaching, a region of 0.5 µm was chosen and applied to either the septum or pole of the cells. Following photobleaching, images were taken every 2 s for 2 min to follow mCherry fluorescence recovery.

The resulting time-lapse images and measured values were further analyzed via Fiji (50) and RStudio (Posit, PBC) using ggplot2 (48) and nlstools (52) packages. The analysis was performed according to the protocol described previously (53). Briefly, the integrated density, area of the region of interest, and mean background fluorescence values were collected for total cell, bleached spot and background. Measured values were used in calculations of the corrected total cell fluorescence (CTCF):

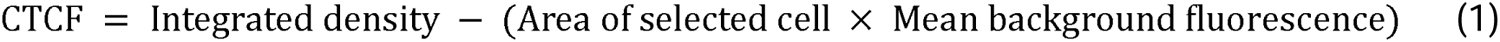

CTCF of the bleached region was divided by the CTCF of the whole cell to account for bleaching during imaging. The normalized recovery values were calculated, and an exponential model was fit to the data:

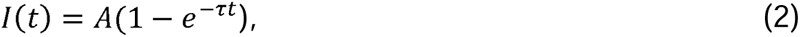

where *I*(*t*) – normalized FRAP curve; *A* – a final value of the recovery; *τ* – fitted parameter; *t* – time after the bleaching event. Half-time of the TatA-mCherry fluorescence recovery (*T*1/2) was determined:

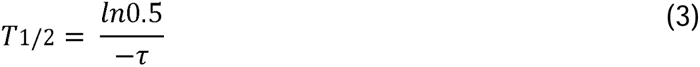

### Single-particle tracking

Overnight cultures of CEK01 and CEK02 strains were diluted to an OD_600_ of 0.5 and grown at 30 °C, 200 rpm until an OD_600_ of ∼2 was reached. 1 mL of each culture was stained with the HaloTag ligand TMR [final concentration of 50 nM] (Promega) at 30 °C, 200 rpm, for 30 min. After staining, cells were harvested at 4,000 rcf, for 3 min and washed five times in sterile filtered TSEMS [50 mM Tris-HCl pH 7.4, 50 mM NaCl, 10 mM EDTA, 0.5 % sucrose filtered through a 0.2 µm filter].

Before imaging, glass slides were cleaned with 1 M KOH, washed with ddH_2_O and dried with pressurized air. Low-melting agarose pads [1 % agarose in sterile filtered TSEMS] (agarose, low gelling temperature, Sigma-Aldrich) were prepared using gene frames (1.0 x 1.0 cm, ThermoFisher Scientific). Cells were mounted on agarose-coated slides and covered with high-performance cover glasses (0.170 ± 0.005 mm thickness).

Images were acquired via an Elyra 7 inverted microscope equipped with two pco.edge sCMOS 4.2 CL HS cameras (PCO AG), connected through DuoLink (Zeiss), only one of which was used in this study. Cells were observed through an alpha Plan-Apochromat 63x/1.46 Oil Korr M27 Var2 objective in combination with an Optovar 1x magnification changer (Zeiss), yielding a pixel size of 97 nm. The focus was automatically maintained via a Definite Focus.2 system (Zeiss). TMR fluorescence was excited with a 561 nm laser with 50% intensity in TIRF mode [62 ° angle], and signals were observed through a multiple beam splitter (405/488/561/641 nm) and laser block filters (405/488/561/641 nm) followed by DuoLink SR DUO filter module (Zeiss; secondary beam splitter: LP560, emission filters: LP570).

For each time-lapse experiment, 15,000 frames were taken with 20 ms exposure time and 4 ms laser transfer time (in total 24 ms).

Cells were selected via Oufti (54) for further analysis. For single-particle tracking, spots were identified via the LoG Detector algorithm in TrackMate v6.0.1 (55) in Fiji 1.53g (50). The estimated spot diameter used for the identification of the fluorescent spots was set to 0.5 µm, and the signal-to-noise ratio threshold was set to 7. The median filter and sub-pixel localization were activated. To limit the detection of ambiguous signal, frames belonging to the TMR bleaching phase (10,000 frames) were removed from the time-lapses prior to the identification of the spots. Tracks were identified via Simple LAP Tracker algorithm, with a maximum linking distance of 300 nm, two frame gaps allowed, and a gap closing distance of 500 nm. Only tracks that persisted for a minimum of 5 frames were used for further analysis (TatA-HaloTag-TMR in CEK01: 4,996 tracks; TatA-HaloTag-TMR in CEK02: 5,740 tracks).

The resulting tracks were subjected to stationary localization (SLA), mean squared displacement (MSD) and square displacement (SQD) analysis in SMTracker 2.0 (56, 57).

The dwell time distribution was determined for a confinement radius of 97 nm and a minimum number of steps within the confinement space of 5 steps. Dwell time distribution was fitted with one and two components, respectively.

The average MSD was calculated for four separate time points per strain [exposure of 20 ms - τ = 24, 48, 72 and 96 ms], followed by fitting a linear equation to the data. The last time point of each track was excluded to avoid track-ending-related artefacts.

The cumulative probability distribution of the SQDs was used to estimate the diffusion constants and relative fraction of up to three diffusive states (fast mobile, slow mobile, and confined). The models were then compared via F-test to verify whether an increase in the number of components could be justified. Further, more complex models were accepted only when a statistically significant F-test (p-value < 0.05) was coupled with a decrease in the associated Bayesian information criterion (BIC) of more than 5 %. The diffusion coefficients and population fraction sizes obtained via the CDF fitting were then used to fit the jump distance probability distributions. Diffusion constants were determined simultaneously for the compared conditions (TatA-HaloTag-TMR in control and Δ*pldP* genomic backgrounds), therefore allowing for a more direct population fraction comparison.

## Results

### PldP is an active ATP-hydrolyzing enzyme

Based on sequence analysis performed using BLAST (58), PldP (Cg1610) is 42.26 % identical to *C. glutamicum* ParA (Cg3427) and 53.44 % identical to *Bacillus subtilis* Soj (Bsu40970). ParA proteins possess a deviant Walker A motif, involved in ATP binding (59, 60). We performed multiple sequence alignments of PldP, ParA, and Soj protein sequences using CLUSTALW (61) and confirmed that PldP contains a conserved deviant Walker A motif (Figure 1A). Compared with the classical Walker A motif, the deviant motif is characterized by a signature lysine at the beginning of the KGGVGKTT amino acid sequence (60, 62). Apart from the Walker A motif, PldP, ParA, and Soj share a conserved aspartate (D74 in PldP; Figure 1A). Previously, this aspartate residue has been shown to be essential for ATP hydrolysis in *Thermus thermophilus* Soj (63). Therefore, based on the presence of two conserved ATP-dependent motifs, we speculated that PldP also possesses ATPase activity. To characterize the hydrolytic activity of PldP, the residues K50 and D74 were mutated to alanine, which was expected to compromise ATP binding and ATP hydrolysis, respectively.

**Figure 1:**
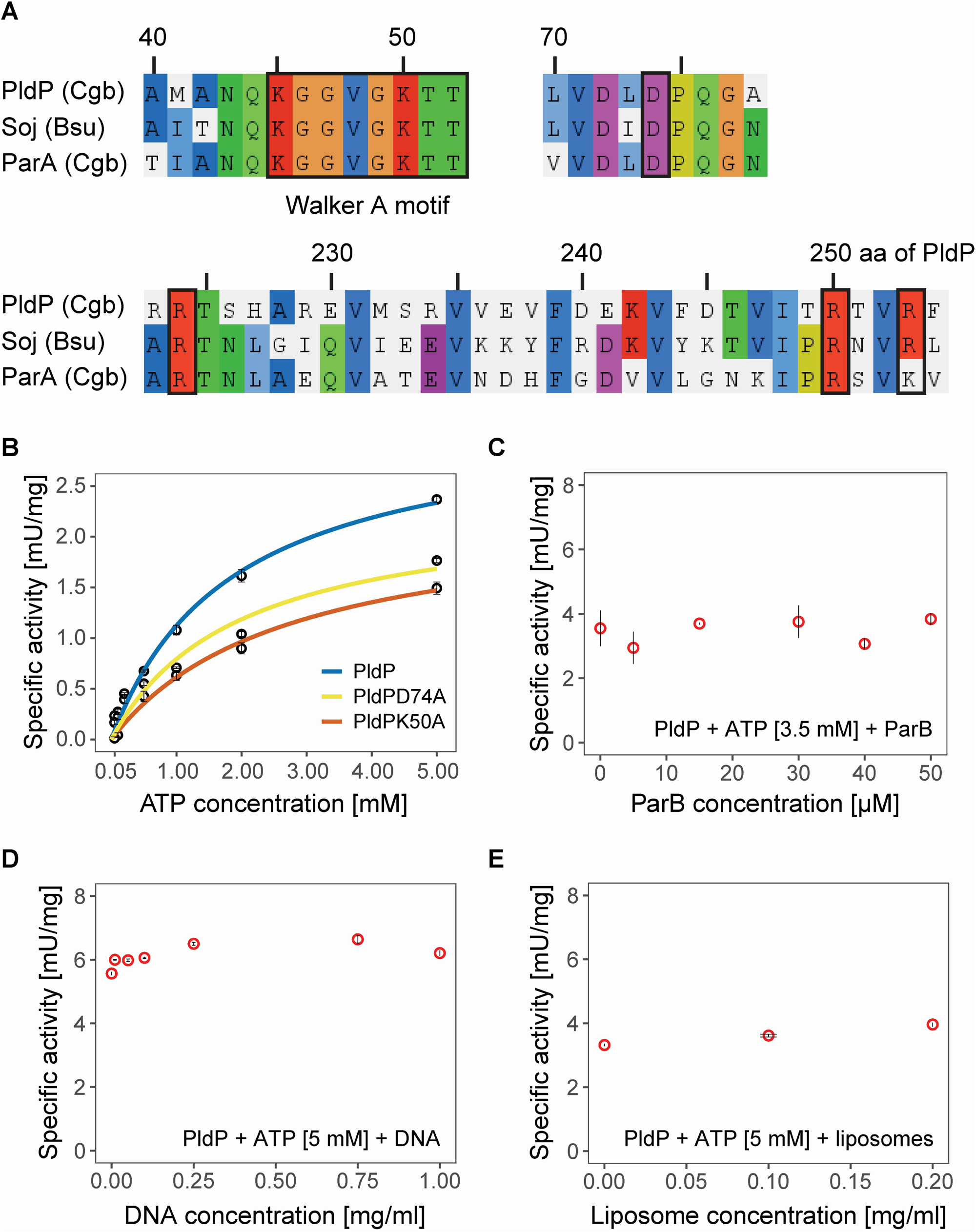
PldP is an ATPase, and its activity is stimulated by the addition of DNA but not ParB or liposomes. **(A)** PldP and ParA from *C. glutamicum* and Soj from *B. subtilis* share conserved Walker A motif, aspartate and arginines. Multiple sequence alignment was performed via CLUSTALW (61) and visualized with AliView (87). Amino acids are counted based on the PldP protein sequence. **(B)** ATPase activities of two PldP variants, K50A and D74A, are decreased. Specific activities of PldP, PldP^K50A^ and PldP^D74A^ follow Michaelis-Menten kinetics. The ATPase activity of PldP does not depend on the presence of ParB **(C)** and liposomes **(E)**; however, it is slightly stimulated by the addition of DNA **(D)**. ATPase activities **(B, D, and E)** were measured using EnzChek^TM^ phosphate assay, where PldP concentration was constant for each setting [5 µM]. In **(C),** the PldP activity in the presence of ParB was determined using the Malachite Green phosphate assay. The measurement of ATPase activity for each sample and condition was performed in three technical replicates. Error bars represent standard deviations. mU – milli-units.

For biochemical assays, hexa-histidine-tagged wild-type PldP and its variants, PldP^K50A^ and PldP^D74A^, were recombinantly produced in *E. coli* BL21(DE3)/pLysS. These proteins were purified via Ni-NTA affinity chromatography (Figure S1), and their activities were tested using EnzChek^TM^ or the Malachite Green phosphate assay. For each replicate, the protein concentration was 5 µM, and ATP concentrations were varied from 0.05 to 5 mM. PldP revealed very low basal ATPase activity that followed Michaelis-Menten kinetics. As expected, the ATPase activities of PldP^K50A^ and PldP^D74A^ were reduced (Figure 1B). The highest maximum rate of ATPase activity (V_max_) was reached by wild-type PldP [V_max_ = 3.20 mU/mg]. PldP^K50A^ ATPase kinetics exhibited the highest K_m_ [K_m_ = 2.70 mM] (Table 1), demonstrating that the intended lowered ATP affinity of this variant was achieved, although the effect was smaller than expected.

**Table 1:** Rates of ATPase activities of PldP and its variants.

| Protein | $V_{\max}$<br>[mU/mg] | $K_m$<br>[mM] |
| --- | --- | --- |
| **PldP** | 3.20 | 1.85 |
| **PldP<sup>K50A</sup>** | 2.27 | 2.70 |
| **PldP<sup>D74A</sup>** | 2.34 | 1.95 |

ParA-like proteins often bind to charged surfaces, such as those provided by DNA or membranes, to function in intracellular transport (13, 17, 64). *B. subtilis* Soj was shown to bind DNA (65, 66) by arginine residues R189 and R218 (66). Multiple sequence alignment confirmed that PldP also contains conserved arginine residues (R224, R250, and R253; Figure 1A). Consequently, we hypothesized that the ATPase activity of PldP may be affected by DNA. PldP activity was tested in the presence of ATP [5 mM] and salmon sperm DNA at varying concentrations [0.01 – 1 mg/mL]. However, PldP ATPase activity was only slightly influenced by the addition of DNA (Figure 1D). We then tested a potential influence of membrane binding, similarly to the lipid-stimulated ATPase activity of MinD (62, 67). Here, we tested PldP ATPase activity in the presence of liposomes [0.1, 0.2 mg/mL]. However, the addition of liposomes did not affect PldP activity (Figure 1E). Finally, the PldP ATPase activity was measured in the presence of corynebacterial ParB. In *Caulobacter crescentus*, the addition of ParB was shown to stimulate ParA ATPase activity in a DNA-dependent manner (17). In *B. subtilis*, Soj ATPase activity increases 70-fold in the presence of Spo0J and DNA (68). We therefore tested the ATP-dependent [3.5 mM] activity of PldP in the presence of ParB [0 – 50 µM]. As a result, we did not observe any influence of ParB on PldP (Figure 1C), indicating that ParB may not act in the same system as PldP, and consequently, the PldP partner protein has yet to be identified.

In summary, we conclude that PldP is an ATPase with low basal ATPase activity that can be only slightly stimulated by DNA and not at all by phospholipid membranes or ParB, suggesting that unknown interaction partners must be involved.

### The cell growth phase determines the spatiotemporal positioning of PldP

Previously, we reported that PldP localizes to the midcell and in patches across the nucleoid (23). Here, we aimed to characterize the localization of PldP further. Accordingly, allelic replacement of *pldP* by *pldP-eYFP* was performed, which allowed for preserving physiological levels of *pldP* gene expression. The fusion of PldP with eYFP was verified using in-gel eYFP fluorescence (Figure S2A). *C. glutamicum* RES167 *pldP::pldP-eYFP* cells (CBK075) were analyzed in the exponential and stationary phases of the cell cycle (Figure 2). We found that prior to cell division, PldP-eYFP localized at the quarter-cell (future midcell) positions and at midcell positions in the exponential phase. A clear midcell positioning was observed in the predominantly short cells of the stationary phase.

**Figure 2:**
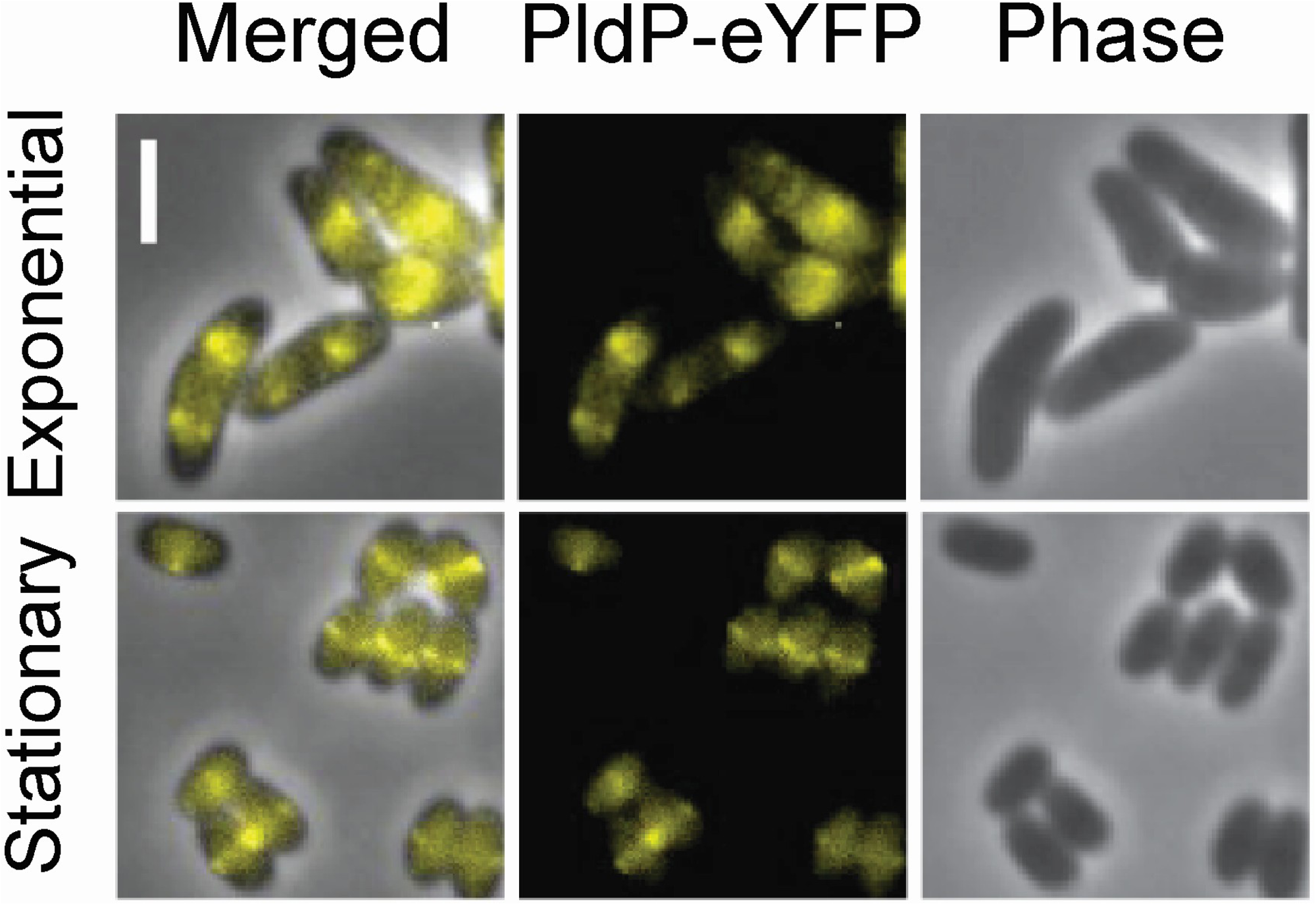
PldP localizes to the cell-quarter positions and septa in a cell cycle-dependent manner. *C. glutamicum* RES167 *pldP::pldP-eYFP* cells (CBK075) were observed in the exponential and stationary phases. PldP-eYFP fluorescence is shown in yellow (false coloured). In the exponential phase cells, PldP-eYFP localizes mostly to the cell quarters. In the stationary phase cells, PldP-eYFP is found in the midcell. Scale bar – 2 µm.

### TatA localization does not depend on PldP

We next aimed at testing the hypothesis that PldP might be the partitioning ATPase that was suspected to drive the dynamics of the Tat translocon in Actinobacteria. The Tat system functions in the transport of folded proteins across the cell membrane (69). In *Streptomyces coelicolor*, components of the Tat system TatABC are highly dynamic (25). In particular, it has been suggested that Tat proteins may be partitioned to each spore during sporulation (25). This implies that a Par-like system may function in partitioning Tat proteins in aerial hyphae formed by members of the genus *Streptomyces*. To investigate the possible similar interaction between PldP and the Tat system in *C. glutamicum*, which, like *Streptomyces* species, belongs to the Actinomycetes, TatA localization was examined. A fusion of TatA and HaloTag was constructed via allelic replacement and verified using in-gel fluorescence (Figure S2B). Localization of TatA-HaloTag was observed in the wild-type and Δ*pldP* backgrounds. For this purpose, *C. glutamicum* RES167 *tatA::tatA-halotag* (CEK01) and *C. glutamicum* RES167 *ΔpldP tatA::tatA-halotag* (CEK02) cells were stained with HaloTag ligand TMR and with DNA-binding dye Hoechst 33342. We found that the localization of TatA-HaloTag-TMR was not affected by the *pldP* deletion (Figure 3). TatA-HaloTag-TMR signals localized to the cell membrane, including the septum, in both strains. The aberrant cell shape phenotype of the Δ*pldP* strain was in accordance with previous findings (23). Notably, in *C. glutamicum,* TatA was not predominantly localized to the cell poles. However, we observed that TatA-HaloTag-TMR foci were dynamic. This finding prompted us to assess the effect of PldP on the dynamic behavior of TatA.

**Figure 3:**
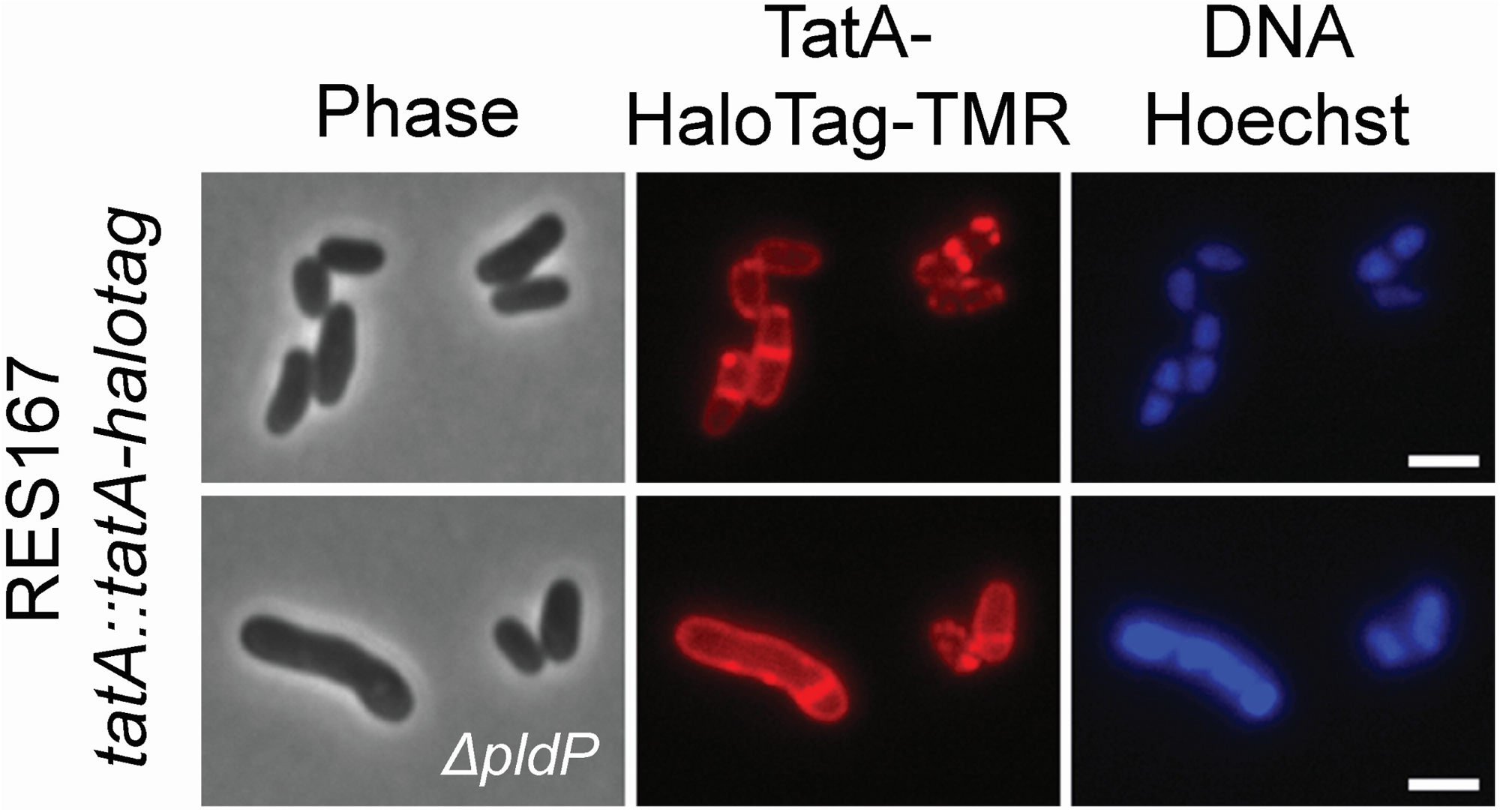
Membrane localization of TatA is not affected by *pldP* deletion. *C. glutamicum* RES167 *tatA::tatA-halotag* (CEK01) and *C. glutamicum* RES167 Δ*pldP tatA::tatA-halotag* (CEK02) cells were stained with HaloTag ligand TMR and DNA-binding dye Hoechst 33342. The cells were observed in the exponential phase. TMR is shown here in red, while DNA stained with Hoechst is shown in blue (false coloured). In both strains, TatA-HaloTag-TMR localizes to the cell membrane, including the septum, and forms rapidly moving clusters. Exposure time: TMR – 50 ms; Hoechst – 40 ms. Scale bar – 2 µm.

### The dynamics of TatA clusters are affected by *pldP* deletion

The mobility of TatA proteins has been shown before in *S. coelicolor* (25) and *E. coli* (70). In *E. coli,* YFP-fused TatA has been found not only as a part of the complex with an average of 25 TatA subunits but also as mobile proteins dispersed at the membrane (70). In case of metallothionein-tagged TatA, average TatA assemblies are composed of 17 +/- 6 TatA protomers, and these clusters are predominantly found at polar or subpolar positions (33), which is where transport takes place (32). Here, we observed the dynamics of TatA clusters via fluorescence recovery after photobleaching (FRAP) experiments. For this purpose, strains expressing *tatA-mCherry* from their native locus were constructed, namely *C. glutamicum* RES167 *tatA::tatA-mCherry* (CAD05) and *C. glutamicum* RES167 Δ*pldP tatA::tatA-mCherry* (CAD06). The fusion of TatA with mCherry was confirmed via Western blotting (Figure S2C) and was fully functional as judged by growth experiments. The mCherry fluorophores in a circular area of 0.5 µm diameter were bleached. The area of bleaching covered either the septum or the cell pole in the exponential phase cells. Following bleaching, the recovery of mCherry fluorescence was recorded by taking images every ∼2 s for the duration of ∼2 min (Figure 4A). We observed that TatA-mCherry fluorescence recovery was generally slower in cells lacking *pldP* (Figure 4B). This was true for the cells with septa; however, there was no significant difference in the recovery half-time in the case of cells without septa. The values of the recovery half-time showed a high standard deviation (Table 2). TatA-mCherry foci often moved out of the focal plane during the recording, which might have contributed to the observed high variation. In summary, we suggest that the deletion of *pldP* affects the speed of TatA-mCherry clusters. They seem to move more slowly, although the dynamic behavior of the clusters is preserved.

**Figure 4:**
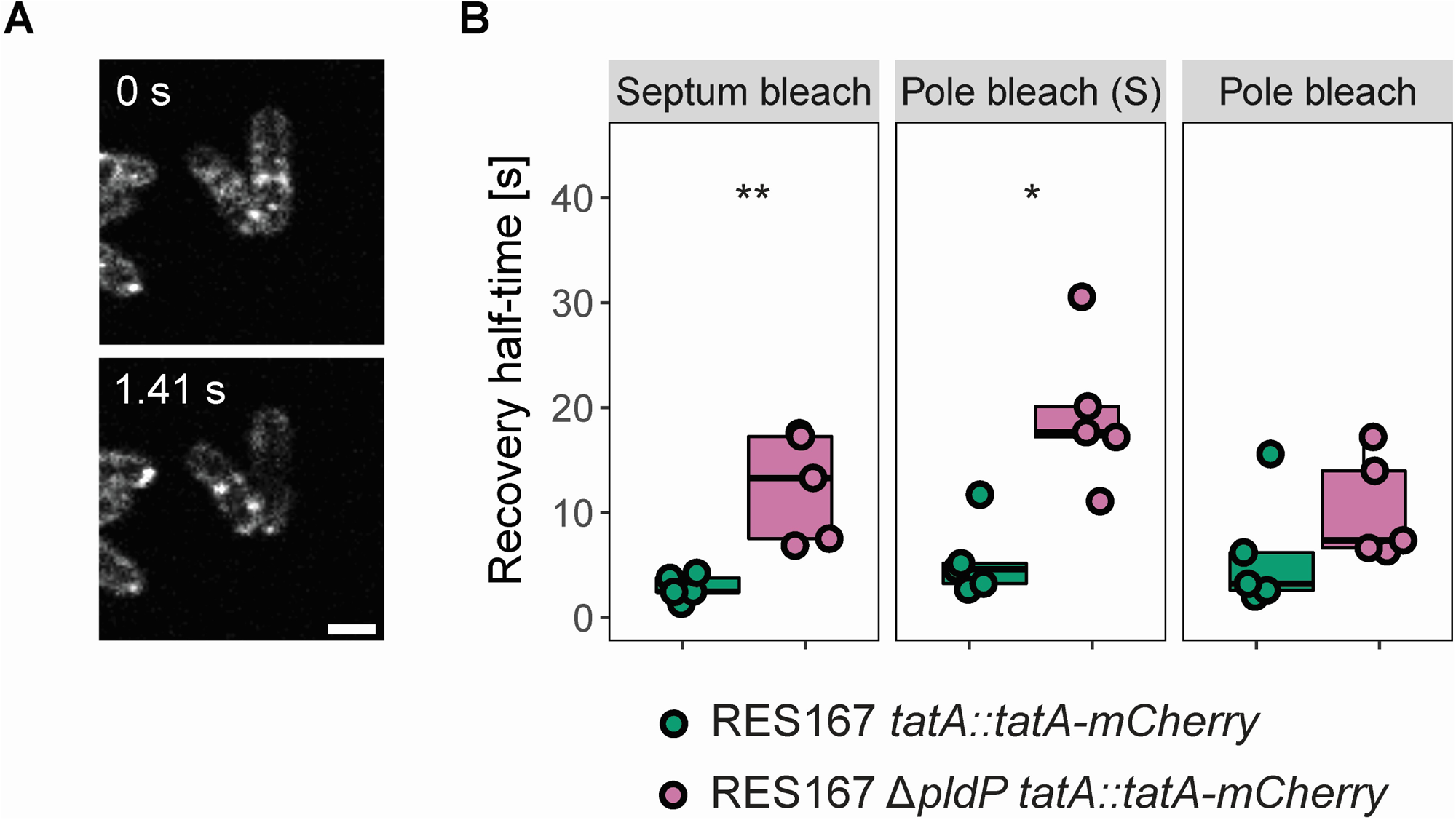
After photobleaching, TatA-mCherry fluorescence recovers more slowly in cells lacking *pldP*. **(A)** *C. glutamicum* RES167 *tatA::tatA-mCherry* (CAD05) and *C. glutamicum* RES167 Δ*pldP tatA::tatA-mCherry* (CAD06) exponential phase cells were used in FRAP experiments. The mCherry fluorophores were bleached in a circular area of 0.5 µm diameter that covered either the septum or the cell pole. Fluorescence recovery was followed by taking images every ∼2 s for the duration of ∼2 min. Fluorescence intensity values were measured and corrected according to the cell area and background fluorescence. **(B)** Fluorescence recovery half-time values are higher in the case of Δ*pldP* cells with septa. * – p < 0.05; ** – p < 0.01 (Mann-Whitney test; number of cells for each condition = 5 cells). S – cells with septa. Scale bar – 2 µm.

**Table 2:**
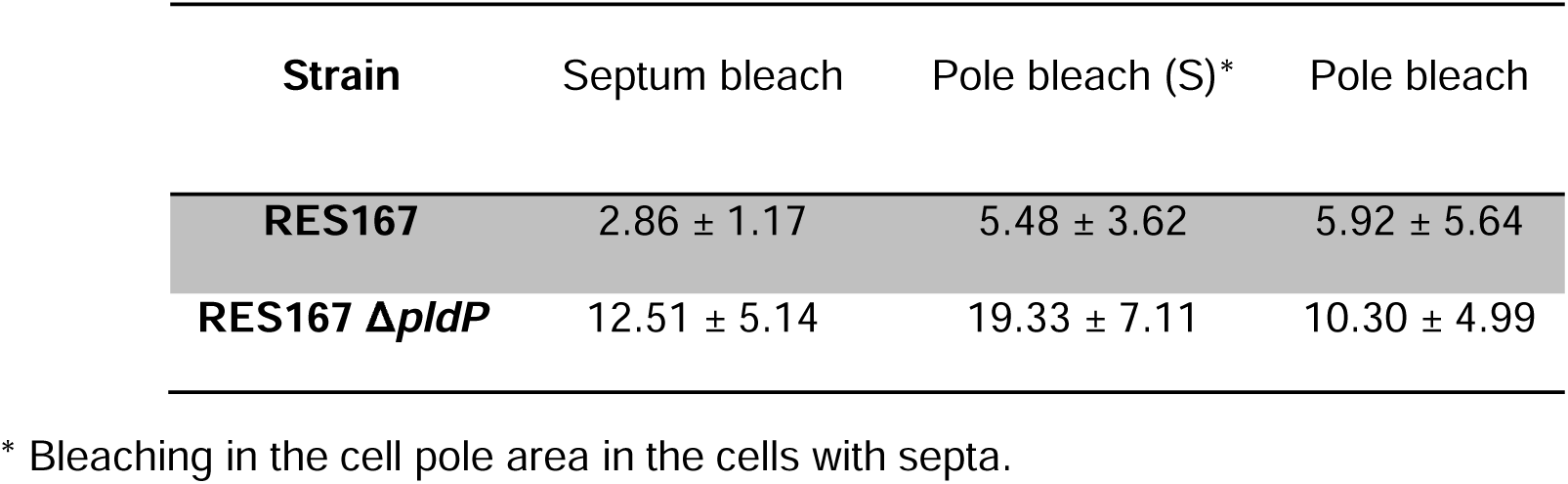
Values of average TatA-mCherry fluorescence recovery half-time ± SD, [s].

To further characterize the impact of PldP on TatA dynamics, we used single-particle tracking (SPT). SPT allows for highly precise analysis of protein dynamics both in space and time. For this specific aim, *C. glutamicum* RES167 *tatA::tatA-halotag* (CEK01) and *C. glutamicum* RES167 Δ*pldP tatA::tatA-halotag* (CEK02) cells were stained with the HaloTag ligand TMR, and the TatA-HaloTag-TMR dynamics were observed. In each time-lapse experiment, a total of 15,000 frames were taken at 20 ms exposure time. Spots and tracks were identified using TrackMate v6.0.1 (55) in Fiji (50) [CEK01 – 4996 tracks; CEK02 – 5740 tracks]. The resulting tracks were analyzed in SMTracker 2.0 (56, 57). We found that TatA-HaloTag-TMR exhibited slower dynamics in the cells lacking *pldP*. The difference was rather subtle, as estimated by mean squared displacement (MSD) analysis [diffusion coefficients: D_CEK01_ = 0.015 µm^2^ s^-1^; D_CEK02_ = 0.010 µm^2^ s^-1^] (Figure 5B). It should be noted that MSD did not account for separate protein populations; therefore, the square displacement (SQD) and jump distance (JD) analysis was preferred (Figure S3). The cumulative distribution function (CDF) of the SQD was used to assess the relative populations of TatA-HaloTag-TMR. The one- and two-component models were fitted to the empirical CDF (ECDF; Figure 5D). The fitting of the three-component model was not justified according to the F-test (see Materials and Methods). Diffusion coefficients for each population were determined simultaneously for two conditions (CEK01 and CEK02 genomic backgrounds) to achieve a direct population comparison. As a result, we observed that TatA-HaloTag-TMR molecules were contained in two relative populations – confined with almost no mobility and slow mobile (Figure 5A). Importantly, the confined population was increased in the strain with *pldP* deletion [increase from 61.8 to 67.5 %]. In other words, it means that TatA-HaloTag-TMR moved more slowly in the absence of PldP. These observations correspond to the FRAP results mentioned above.

**Figure 5:**
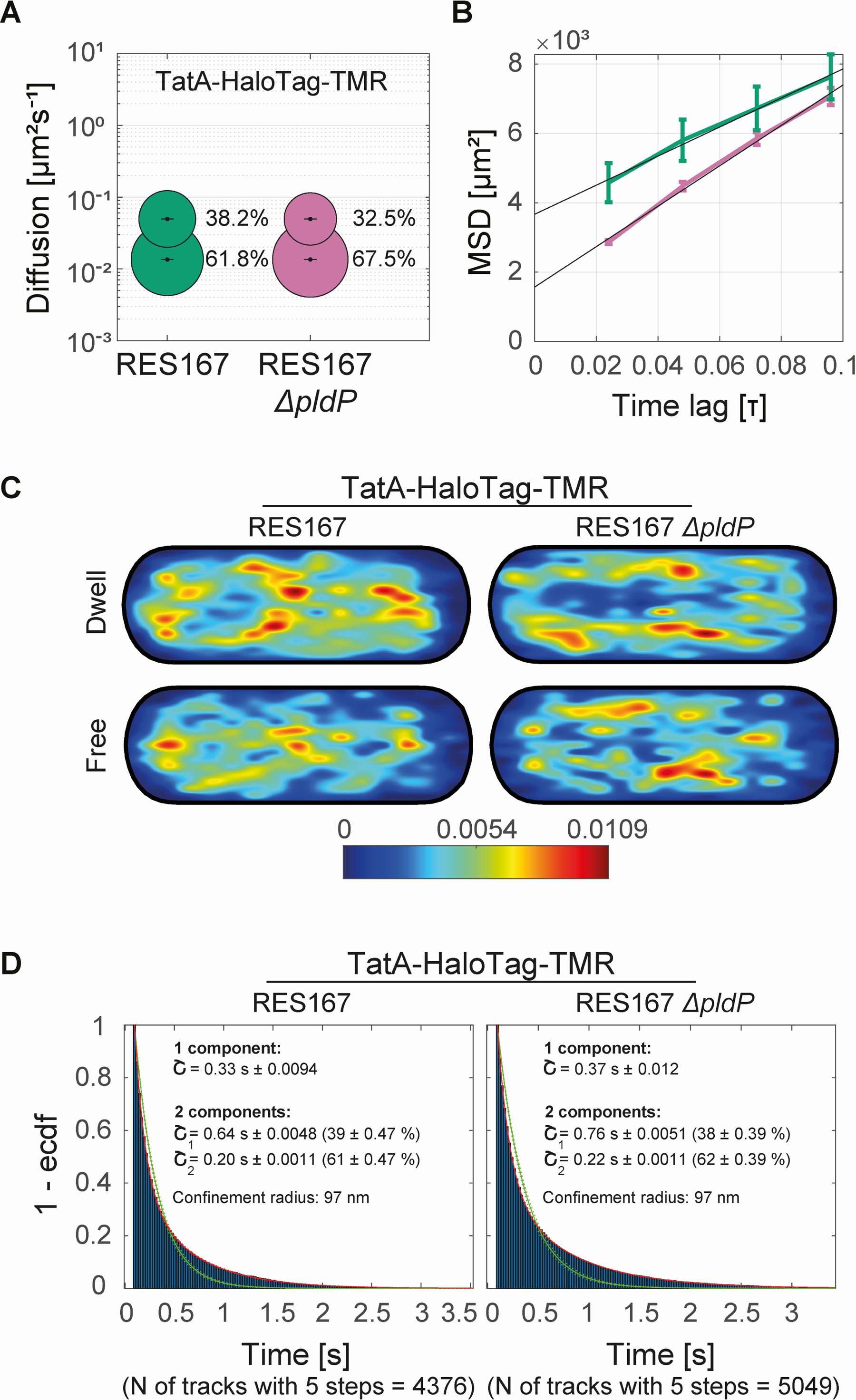
The deletion of *pldP* leads to an increase in TatA confined population. *C. glutamicum* RES167 *tatA::tatA-halotag* (CEK01) and *C. glutamicum* RES167 Δ*pldP tatA::tatA-halotag* (CEK02) cells were used for SPT experiments. **(A)** The cumulative distribution function (CDF) of the square displacements (SQD) was used to estimate the diffusion coefficients and relative populations of TatA-HaloTag-TMR of up to two diffusive states. Here, the diffusion coefficients and corresponding TatA-HaloTag-TMR populations are shown in the bubble plot. TatA-HaloTag-TMR molecules are contained in two populations – confined (lower diffusion rates) and slow mobile (higher diffusion rates). The confined population is increased upon *pldP* deletion [increase from 61.8 to 67.5 %]. **(B)** The plot of the mean squared displacements (MSD) highlights the differences in TatA-HaloTag-TMR diffusion between CEK01 (green) and CEK02 (purple). The average MSD was calculated for four separate time points per strain, and the linear regression model was fitted to the data. **(C)** Confinement heatmaps show the localization of dwelling and free TatA-HaloTag-TMR molecules in an averaged cell. The probability of the presence of tracks is shown as a spectrum from blue to red (from the lowest to highest probability, respectively). In CEK01 cells, TatA-HaloTag-TMR molecules localize to the cell membrane, including the septum. In the case of CEK02 cells, the septal localization of TatA-HaloTag-TMR is not evident because the septum does not form precisely in the midcell. **(D)** Empirical cumulative distribution function (ECDF) with fitted one- and two-component models. Analysis was performed using SMTracker v2.0 (56, 57).

In agreement with the reported findings, we confirmed the localization of TatA-HaloTag-TMR at the cell membrane in both strains (Figure 5C). The septal localization was evident only for CEK01 cells and not for CEK02 (Δ*pldP*). This is explained by the fact that the septum does not form precisely in the middle of the cell with *pldP* deletion (Figure 3). Therefore, in CEK02 (Δ*pldP*), the septal localization of TatA-HaloTag-TMR appears dispersed throughout the cell length. Overall, with FRAP and SPT data, we show that *pldP* deletion led to a decrease in the dynamics of TatA proteins, which were present in confined and slow mobile populations.

### The deletion of *pldP* does not influence Tat-mediated protein secretion

As the mobility of TatA was altered in the absence of PldP, we addressed the potential effect of the *pldP* deletion on Tat transport. For that purpose, we analyzed the transport of two Tat-dependently secreted proteins of *C. glutamicum*, the alkaline phosphatase PhoD (71) and a Rv2525c-like glycoside hydrolase-like domain-containing protein Cg0955 in the reference genome of *C. glutamicum* ATCC 13032 (Bielefeld) (72). Based on the amino acid sequence alignment via BLAST (58), Cg0955 was 48.97 % identical to the peptidoglycan hydrolase Rv2525c from *Mycobacterium tuberculosis* (73, 74). Notably, when we started this analysis, the Tat-specificity of the signal peptide from Cg0955 was only predicted and not verified. It has been previously shown to be the most efficient signal peptide in *C. glutamicum* R (therein CgR0949, gene locus CGR_RS04950), resulting in 150-fold increased amounts of secreted α-amylase compared to the secretory protein PS2 (75).

As shown in Figure 6, twin-arginine-specific transport could be demonstrated for both PhoD and Cg0955, as the original RR signal peptide resulted in extracellular, mature forms of the proteins (lanes E of RR constructs), whereas the RR>KK mutated signal peptides did not promote transport (no signal in lanes E in case of KK constructs). Note that the charge-conservative RR>KK exchange specifically abolishes Tat transport (76). While the KK-precursor of Cg0955 accumulated in the cytoplasm, the non-transported KK-PhoD was largely degraded, and only the membrane-associated precursor was unaffected, indicating an instability of PhoD in the cytoplasmic compartment when transport is blocked. Importantly, the deletion of *pldP* had no significant impact on Tat transport in both cases (compare lanes WT with Δ*pldP* in both blots). Therefore, we conclude that PldP has no influence on active Tat-dependent secretion, despite the above-described effects on TatA dynamics.

**Figure 6:**
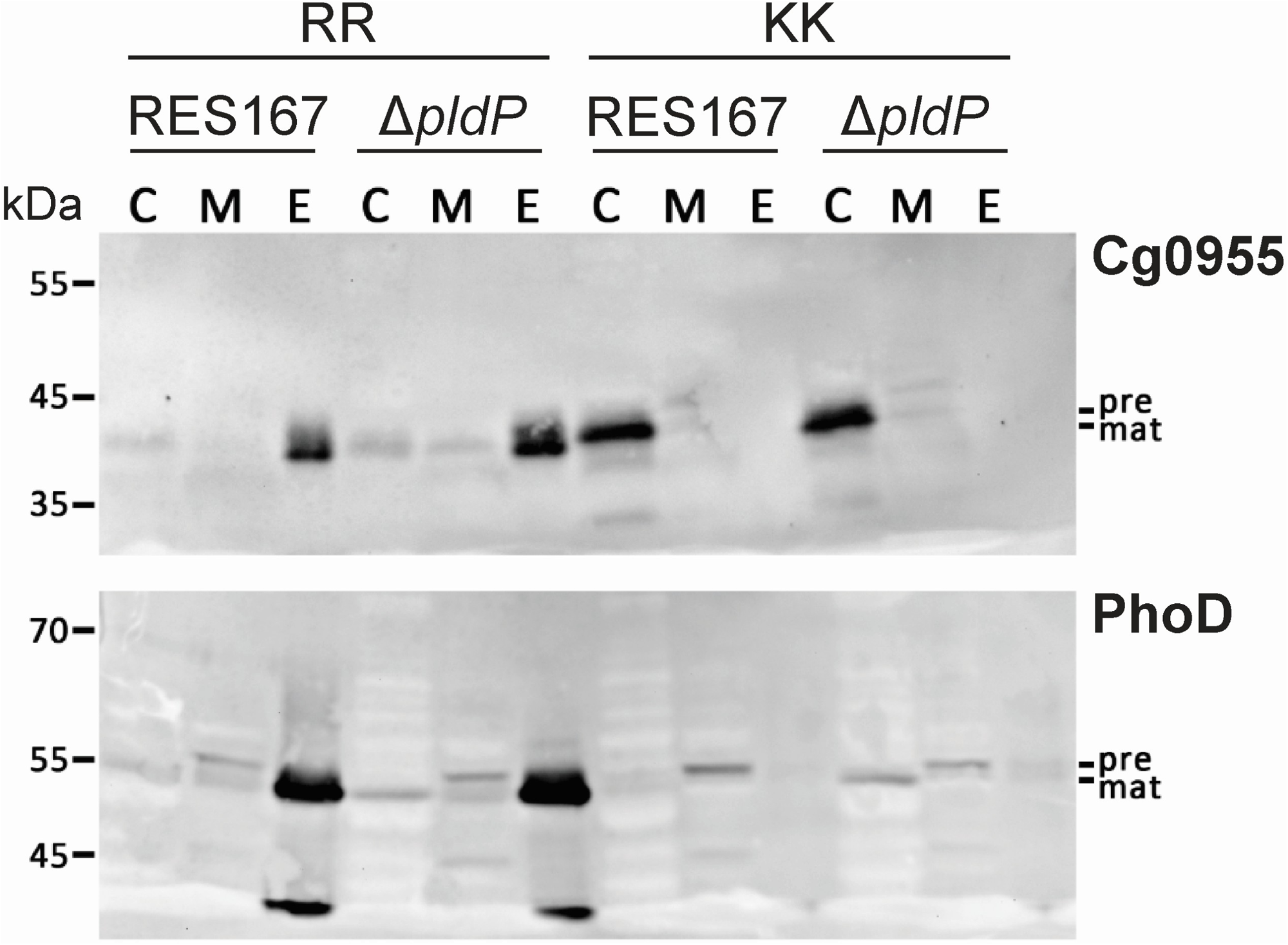
The deletion of *pldP* does not influence Tat-mediated secretion. Transport was analyzed by SDS-PAGE/Western blotting of cytoplasmic (C), membrane (M), and extracellular (E) fractions of *C. glutamicum* RES167 and its Δ*pldP* mutant (CDMB01-08), harboring plasmids for constitutive expression of Tat substrates Cg0955 (upper blot) and PhoD (lower blot) with their native RR-signal peptides (RR) or their RR>KK mutated variants (KK). The upper blot shows the results with pCLTON1-*P_sod_-cg0955-H6* (pDMB-RR-0955-H6; RR-lanes) and pCLTON1-*P_sod_-KK-cg0955-H6* (pDMB-KK-0955-H6; KK-lanes), whereas the lower blot shows the results obtained with pCLTON1-*P_sod_-phoD-H6* (pDMB-RR-PhoD-H6; RR-lanes) and pCLTON1-*P_sod_-KK-phoD-H6* (pDMB-KK-PhoD-H6; KK-lanes). Tat-mediated transport is demonstrated for both Cg0955 and PhoD with the original RR signal peptides (lanes E of RR constructs). The mutation RR>KK in the signal peptide did not promote transport (lanes E of KK constructs). The KK-precursor of Cg0955 accumulated in the cytoplasm, while the cytoplasmic KK-precursor of PhoD was mainly degraded, and only the membrane-associated PhoD precursor was unaffected. The tags were detected using hexahistidine-specific antibodies. The extracellular fractions were enriched by Ni-NTA affinity chromatography to an equivalent volume. Positions of protein markers are indicated on the left, and those of precursor (pre) and mature proteins (mat) on the right.

### The deletion of *pldP* results in spatial mislocalization of Tat-mediated protein transport

One of the consequences of the decreased TatA dynamics could be altered spatial localization of Tat-mediated protein secretion. To observe the influence of PldP on the localization of Tat secretion, first, we generated a mNeonGreen fused to a Tat secretion signal of the hypothetical glycoside hydrolase Cg0955 (Figure S4A). The overexpression of *tatSP-mNeonGreen* in *C*. *glutamicum* RES167 pEKEx2-*tatSP-mNeonGreen* (CEK03) and *C. glutamicum* RES167 Δ*pldP* pEKEx2-*tatSP-mNeonGreen* (CEK04) cells was induced by IPTG [10 µM]. After induction, cells were stained with the cell membrane dye FM^TM^4-64. We observed that the mNeonGreen signal localized prominently to the septum in both strains, suggesting that Tat transport takes place near the septum. In the strain that contained PldP, mNeonGreen was also located at the cell envelope regions neighboring the septum (Figure S4B). Generally, the mNeonGreen fluorescence was more uniformly distributed in cells lacking PldP, and fluorescence was dimmer at the cell poles (Figure 7A) and septa (Figure 7B). In summary, these data suggest that a *pldP* deletion results in a more homogeneous targeting of the TatSP-mNeonGreen fusion, and therefore, PldP clearly has an influence on the subcellular localization of Tat-mediated secretion, in line with the observed differences in localization of TatA in *pldP* mutant strains.

**Figure 7:**
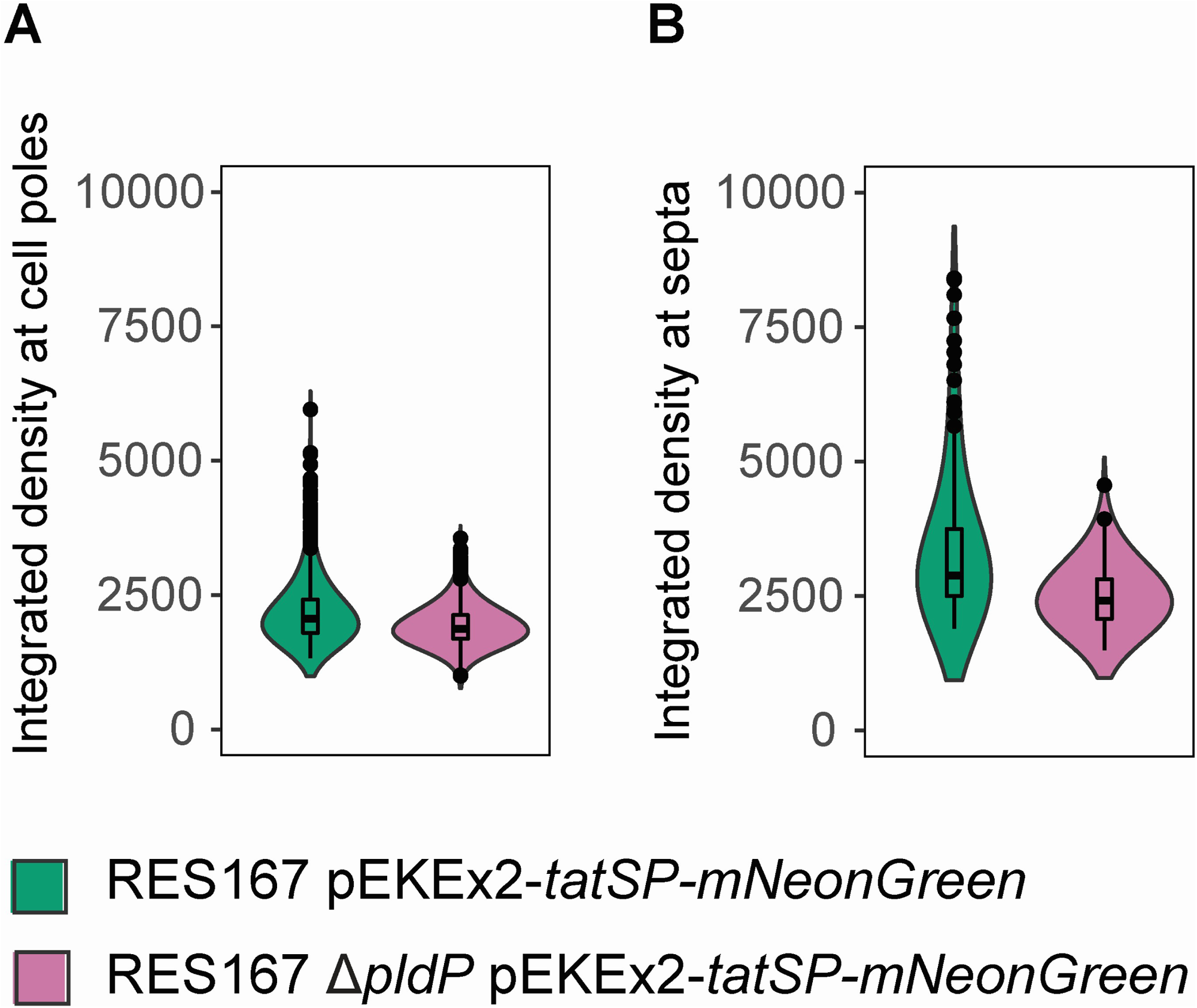
The deletion of *pldP* leads to a decrease in mNeonGreen fluorescence at the cell poles and septa. The overexpression of *tatSP-mNeonGreen* was induced by IPTG [10 µM] in the *C. glutamicum* RES167 pEKEx2-*tatSP-mNeonGreen* (CEK03) and *C. glutamicum* RES167 Δ*pldP* pEKEx2-*tatSP-mNeonGreen* (CEK04) cells. Integrated densities of mNeonGreen were calculated in Fiji (as a product of area and mean grey value of pixels) (50). Integrated density was preferred to fluorescence intensity because the former accounted for the ROI size. The difference in integrated densities between the two strains in each condition is significant (p < 0.0001, Mann-Whitney test; number for each strain and condition: A – 700 cell poles (350 cells); B – 102 septa (102 cells)).

## Discussion

ParA-like proteins are known to localize various protein targets within bacterial cells (10). The actinobacterium *C. glutamicum* harbors an orphan ParA-like protein that was termed PldP, for ParA-like division protein. Deletion of the *pldP* gene results in cell elongation and an increase in asymmetric septum position, resulting in daughter cells of variable cell length (23). Localization of PldP was shown to be dispersed across the nucleoid and enriched at septa. However, the precise function of PldP remained unclear and therefore also the molecular reason for the observed morphology phenotypes associated with a *pldP* null allele. Among those proteins that were speculated to have an active system involved in subcellular dynamics and positioning is the twin-arginine translocation system (Tat) from the actinobacterium *S. coelicolor* (25). Importantly, Tat-mediated protein transport is required for several proteins involved in cell wall metabolism, such as murein hydrolases (31, 77, 78). Loss of these proteins results in similar phenotypes as observed for the *pldP* deletion mutant in *C. glutamicum*. We therefore wanted to explore the possibility that PldP might be involved in the spatial organization of the Tat system in *C. glutamicum*. ParA-like proteins require an active ATP hydrolysis cycle for their positioning activity. Therefore, we first isolated PldP and tested the purified protein for its ATPase activity. As expected, we could confirm a basal ATPase activity of PldP. The ATPase activity is usually enhanced by a binding partner of the ParA-like protein, and we thus tested known interaction partners of ParA-like proteins for their potential effect on PldP. Canonical ParA is triggered by the ParB protein, and MinD proteins are activated by membrane binding and, in the case of *E. coli* MinD, also by a partner protein, MinE. PldP ATPase activity is slightly but significantly stimulated by DNA, suggesting that PldP can bind to DNA. This would be in line with the observed localization of PldP across the nucleoid *in vivo*. In contrast, we did not detect any effect of ParB or membrane vesicle addition. Based on these results, we hypothesize that PldP likely non-specifically binds to DNA, which influences ATP hydrolysis activity, which could account for the dynamic behavior of PldP *in vivo*, which agrees with earlier studies on other ParA-like proteins (17, 20, 21, 63). It is thus very likely that a yet-to-be-identified protein partner further stimulates PldP activity. Mutational analysis in which we substituted the conserved lysine and aspartate residues in the Walker A/switch I motif revealed a significant decrease in the ATPase activity. Often, the substitution of these conserved residues leads to near-complete abolishment of the nucleotide hydrolysis activity. However, we still observed considerable activity for both mutant proteins. Similar behavior has been described for the related ParF protein, in which the corresponding K15Q mutant displayed a reduced activity that could be alleviated by higher ATP concentrations (79).

ATPase activity is likely needed for the observed cell cycle-dependent localization of PldP which includes enrichment at the division plane during cytokinesis. The division septum is not only a place where new cell wall synthesis is initiated to allow daughter cell separation, but it is also a place of increased protein transport. We, therefore, tested a possible involvement of PldP in the dynamics of the Tat translocon. We generated a functional translational fusion to the TatA protein. TatA is the Tat component responsible for the permeabilization of the bacterial membrane (33, 80). It has been shown in *E. coli* that TatA forms larger clusters with an average of ca. 17-25 protomers (70, 81, 82). Based on single-molecule measurements, it was hypothesized that the formation of TatA assemblies starts from tetrameric building blocks (70).

We show here that in *C. glutamicum,* TatA foci localization at cell poles and septa is significantly decreased in cells lacking PldP. While it is impossible to judge differences in spatial localization when observing dynamic clusters, our single-molecule microscopy approach clearly reveals that, on average, TatA clusters are located less frequently at poles and septa in Δ*pldP* strains. SPT data furthermore show that deletion of *pldP* leads to a decrease in TatA mobility, suggesting that PldP contributes to the spatial organization of the Tat components. A potential direct interaction of PldP with TatA could not be demonstrated. Two hybrid experiments did not reveal interaction, and co-localization studies of PldP and TatA were inconclusive, mainly because not all TatA molecules are part of clusters, hence blurring a co-localization analysis. The diffuse presence of smaller TatA assemblies that are not part of larger TatA clusters has also been described in other bacteria (83, 84). However, our findings align well with data from related bacteria, such as *S. coelicolor*, where dynamic positioning of the Tat machinery has been observed (25).

The reduction in TatA dynamics in the absence of PldP did not affect Tat transport of two model substrates, showing that PldP is mechanistically not required for Tat transport. As the reduced TatA dynamics in the absence of PldP already pointed to possible defects in the spatial organization of the transport, we considered that the proper function of transported Tat substrates may depend on a spatially organized transport, i.e. a transport at a specifically required subcellular position. To address this in more detail, we localized the secretion of the fluorescent protein mNeonGreen fused to a Tat-signal peptide. In wild-type and Δ*pldP* mutant cells, the Tat-dependent mNeonGreen was readily observed around the cell periphery, but mNeonGreen transport was most prominent at the septa. We assume that the enrichment of translocated Tat substrates at septa is a result of the specific cell wall architecture. The inward-growing peptidoglycan is still connected to the circumferential cell wall layer, generating the typical pi-shaped structure, observed in transmission EM images (85, 86). Likely, this pi-structure serves as a diffusion barrier, leading to the enrichment of those Tat substrates that are transported at the septum. mNeonGreen quantities at the septum were significantly higher in wild-type compared to Δ*pldP* mutant cells. Since the Tat system is not impaired in its translocation activity in the absence of PldP, this data is best explained by a spatially altered secretion. In other words, if the TatA complex is not efficiently positioned at poles and septa, the correct localization of targeting is compromised, and this is likely the reason for the observed phenotypes.

In conclusion, our study positions PldP as a significant player in the orchestration of cellular processes in *C. glutamicum*. While its exact mechanisms remain to be fully elucidated, the gained insights underscore the critical interplay between PldP ATPases and the cellular dynamics of the Tat translocon. Future research should focus on unravelling the specific interactions through which PldP influences Tat component dynamics. This will contribute to a deeper understanding of bacterial cell biology and the roles of ParA ATPases in maintaining cellular integrity and functionality.

## Acknowledgments

Work in the Bramkamp laboratory is funded by the Deutsche Forschungsgemeinschaft (DFG, BR2915/10-1). The authors thank all members of the Bramkamp lab for critical discussions and comments on the manuscript. We thank Dr. Urśka Repnik from the Central Microscopy facility (Kiel University) for help with the FRAP experiments.

