## Supplemental Material for "The ParA-like ATPase PldP influences the TatA dynamics in *Corynebacterium glutamicum*"

### Supplementary Material

**Table S1: Oligonucleotides used in this study.**

Site-mutagenesis sites are highlighted in yellow; restriction sites (RS) and adjacent binding nucleotides are underlined.

| # | Name | Sequence 5' – 3' | RS | Reference |
|---|---|---|---|---|
| DA001 | PldPK50A_F | GGCGAATCAAGCTGGTGGCGTTG | / | This study |
| DA002 | PldPK50A_R | ATGGCAATGATCGTTGCTG | / | This study |
| DA003 | PldPD74A_F | CGTTGACTTGGCTCCGCAAGGTG | / | This study |
| DA004 | PldPD74A_R | AGCAGGACTTTACGTCCC | / | This study |
| DA005 | pET16b_seq_F | ATCCGGATATAGTTCCTC | / | This study |
| DA006 | pET16b_seq_R | GAATTGTGAGCGGATAAC | / | This study |
| KB001 | ParB_F | CAGCATATGGCTCAGAACAAGGGTTCC | NdeI | This study |
| KB002 | ParB_R | CAGCTCGAGTTATTGGCCCTGGATCAA | XhoI | This study |
| KB003 | PldP_up_F | CATCCTGCAGGAGTGAGTGATGCAGGGAA | SbfI | This study |
| KB004 | PldP_up_R | CATTCTAGAGTCGTTGACGCGGCTGATAA | XbaI | This study |
| KB005 | eYFP_R | CATCCCGGGTTACTTGTACAGCTCGTCCA | XmaI | This study |
| KB006 | PldP_down_F | CATCCCGGGTAGGTTGTTTTTCTA | XmaI | This study |
| KB007 | PldP_down_R | CATGAATTCCGCGGGAGCAGGCGA | EcoRI | This study |
| CD001 | ΔpldP_1_F | CAGAAGCTTAGGTGTATGACAGGGAAA | HindIII | (1) |
| CD002 | ΔpldP_1_R | GGAATGGAGTATGGAAGTTGGAGTCAAACCTTCTTCCTT | / | (1) |
| CD003 | ΔpldP_2_F | CCAACTTCCATACTCCATTCCCCACGCCTCCTTGTGCGG | / | (1) |
| CD004 | ΔpldP_2_R | CAGTCTAGACGCGGGAGCAGGCGAGCT | XbaI | (1) |
| EK43 | TatAup_F | CTGACCTGCAGGTGCATAAAGCAAAAGGCTTTTA | SbfI | This study |
| EK44 | TatAup_R | CGGACTAGTTTCTGGGCCTGGCAGAGAGGTGCGGTTTGGATC | SpeI | This study |
| EK45 | Halo_F | GGAACTAGTATGGCAGAAATCGGTACTG | SpeI | This study |
| EK46 | Halo_R | CAGCAGGCGGCCGCTTAGCCGGAAATCTCGAGC | NotI | This study |
| EK47 | TatAdown_F | ATAAGAATGCGGCCGCAGTTGGGCAGTTTGCATCT | NotI | This study |
| EK48 | TatAdown_R | GCGGGAATTCGCCCCACAGCTGGCT | EcoRI | This study |
| DA007 | -189bp_TatA_F | CTATGACCATGATTACGCCAGATCTTCGGAAATGCATAAAGC | / | This study |
| DA008 | -189bp_TatA_R | TTCTGGGCCTGGCAGTTCTGGGCCTGGCAGAGAGGTGCGGT TTGGATC | / | This study |
| DA009 | mCherry_F | CTGCCAGGCCCAGAACTGCCAGGCCCAGAAATGGTGAGCAA GGGCGAG | / | This study |
| DA010 | mCherry_R | CTGCCCAACTTTACTTGTACAGCTCGTCCATG | / | This study |
| DA011 | TatA+500_F | GTACAAGTAAAGTTGGGCAGTTTGCATCTAAAAAATAAAGTCA TC | / | This study |
| DA012 | TatA+500_R | GAGCTCGGTACCCGGGGATCGCCCCACAGCTGGCTCAG | / | This study |
| DA013 | DA_pK19mobsacB_F | GATCCCCGGGTACCGAGC | / | This study |
| DA014 | DA_pK19mobsacB_R | TGGCGTAATCATGGTCATAGCTG | / | This study |
| GAA57 | pK19_seq_F | GCTTCCGGCTCGTATGTTG | / | This study |
| GAA58 | pK19_seq_R | GCTGCAAGGCGATTAAGTTG | / | This study |

| | | | | |
|---|---|---|---|---|
| EK39 | TatSP_F | ACGCGTCGACTTTGGGAGAACTTTTGGACAT | SalI | This study |
| EK40 | TatSP_R | CGGACTAGTGGCGTTGGCCTTTGGC | SpeI | This study |
| EK41 | mNeonGreen_F | GGAACTAGTATGGTGAGCAAGGGCG | SpeI | This study |
| EK42 | mNeonGreen_R | TCGCGGATCCTTACTTGTACAGCTCGTC | BamHI | This study |
| DMB001 | Cg-Psod-AatII-F | GATCGACGTCAACAAGCTGCTCGAGGGAAATC | AatII | This study |
| DMB002 | Cg-Psod-AvrII-R | GCGCCCTAGGAAAATCCTTTCGTAGGTTTCCGCACCGAGCATATACATC | AvrII | This study |
| DMB003 | Cg-cg0955-AvrII-F | TATATCCTAGGATGCAAATAAACCGCCGAGGCTTCTTAAAAGCCAC | AvrII | This study |
| DMB004 | Cg-cg0955-BamHI-R | TATATGGATCCGCTAGATAGGGTGCCAAGAATCTTGATTACTTGGTTGATG | BamHI | This study |
| DMB005 | AvrII-removal | GATACCCCCGCTAGGTATCGGACAC | / | This study |
| DMB006 | Cg-phoD-AvrII-F | TATATCCTAGGATGCCACAGTTAAGCAGACGCCAG | AvrII | This study |
| DMB007 | Cg-phoD-BamHI_R | TATATGGATCCAGTAGTGAATCCGACGCCTG | BamHI | This study |
| DMB008 | Cg-cg0955-KK-F | GATTTTCCTAGGATGCAAATAAACAAAAAGGGCTTCTTAAAAGCCACCACAGGAC | AvrII | This study |
| DMB009 | Cg-phoD-KK-F-dm | CCTAGGATGCCACAGTTAAGCAAAAAGCAGTTCTTGCAGACAACCGCCG | AvrII | This study |

**Table S2: Plasmids used in this study.**

| Name | Description | Reference |
|---|---|---|
| pET16b | *E. coli* inducible heterologous expression vector, Amp$^R$, 10 x N-terminal histidines (His-Tag) followed by Factor Xa site, *lacI* | Novagen |
| pAS001 | pET16b-*pldP* | This study |
| pDA001 | pET16b-$pldP^{K50A}$ | This study |
| pDA002 | pET16b-$pldP^{D74A}$ | This study |
| pKB001 | pET16b-*parB* | This study |
| pK19*mobsacB* | *C. glutamicum* mobilization vector for allelic exchange, Km$^R$, pK18 *oriV*, *sacB*, *lacZα* | (2) |
| pKB002 | pK19mobsacB-*pldP-eYFP* | This study |
| pCD121 | pK19mobsacB-Δ*pldP* | (1) |
| pEK01 | pK19mobsacB-*tatA-halotag* | This study |
| pDA003 | pK19mobsacB-*tatA-mCherry* | This study |
| pUC19 | *E. coli* pUC cloning vector, $P_{lac}$, Amp$^R$ | (3) |
| pCLTON1 | *C. glutamicum* tetracycline-inducible expression vector, *tetR* under control of $P_{gap}$ promoter, Km$^R$ | (4) |
| pSG1154 | C-terminal fusion vector, *bla amyE3'* Spc$^R$ $P_{xyl}$-'*gfpmut1 amyE5'* | (5) |
| pDMB-RR-0955-H6 | pCLTON1-$P_{sod}$-*cg0955-H6* | This study |
| pDMB-KK-0955-H6 | pCLTON1-$P_{sod}$-*KK-cg0955-H6* | This study |
| pDMB-RR-PhoD-H6 | pCLTON1-$P_{sod}$-*phoD-H6* | This study |

| | | |
|---|---|---|
| pDMB-KK-PhoD-H6 | pCLTON1-$P_{sod}$-*KK-phoD-H6* | This study |
| pEKEx2 | *E. coli* and *C. glutamicum* shuttle vector for IPTG-inducible gene expression, $Km^R$, *lacIq*, *tac* promoter, pUC18 multiple cloning site | (6) |
| pEK02 | pEKEx2-*tatSP-mNeonGreen* | This study |

**Table S3: Bacterial strains used in this study.**

| # | Description | Genotype | Reference |
|---|---|---|---|
| | *Escherichia coli* | | |
| / | *E. coli* NEB® 5-alpha | *fhuA2Δ(argF-lacZ)U169 phoA glnV44 Φ80Δ(lacZ)M15 gyrA96 recA1 relA1 endA1 thi-1 hsdR17* | NEB |
| EAD04 | *E. coli* NEB® 5-alpha pET16b-*pldP* | NEB® 5-alpha with IPTG-inducible pET16b-*pldP* | This study |
| EAD05 | *E. coli* NEB® 5-alpha pET16b-$pldP^{K50A}$ | NEB® 5-alpha with IPTG-inducible pET16b-$pldP^{K50A}$ | This study |
| EAD06 | *E. coli* NEB® 5-alpha pET16b-$pldP^{D74A}$ | NEB® 5-alpha with IPTG-inducible pET16b-$pldP^{D74A}$ | This study |
| EEK01 | *E. coli* NEB® 5-alpha pK19*mobsacB-tatA-halotag* | NEB® 5-alpha with pK19*mobsacB-tatA-halotag* | This study |
| EAD01 | *E. coli* NEB® 5-alpha pK19*mobsacB-tatA-mCherry* | NEB® 5-alpha with pK19*mobsacB-tatA-mCherry* | This study |
| EEK02 | *E. coli* NEB® 5-alpha pEKEx2-*tatSP-mNeonGreen* | NEB® 5-alpha with IPTG-inducible pEKEx2-*tatSP-mNeonGreen* | This study |
| / | *E. coli* BL21(DE3)pLysS | F- *ompT hsdS*B ($r_B^- m_B^-$) *gal dcm* (DE3) pLysS ($Cam^R$) | ThermoFisher Scientific |
| EAD18 | *E. coli* BL21(DE3)pLysS pET16b-*pldP* | BL21(DE3)pLysS with IPTG-inducible pET16b-*pldP* | This study |
| EAD19 | *E. coli* BL21(DE3)pLysS pET16b-$pldP^{K50A}$ | BL21(DE3)pLysS with IPTG-inducible pET16b-$pldP^{K50A}$ | This study |
| EAD20 | *E. coli* BL21(DE3)pLysS pET16b-$pldP^{D74A}$ | BL21(DE3)pLysS with IPTG-inducible pET16b-$pldP^{D74A}$ | This study |
| EPH04 | *E. coli* BL21(DE3)pLysS pET16b-*parB* | BL21(DE3)pLysS with IPTG-inducible pET16b-*parB* | This study |
| | *Corynebacterium glutamicum* | | |
| RES167 | *C. glutamicum* RES167 | Restriction-deficient mutant, otherwise considered wild-type (ATCC 13032 *ΔmluI*) | (7) |
| CBK075 | *C. glutamicum* RES167 *pldP::pldP-eYFP* | RES167 *pldP::pldP-eYFP* | This study |
| CDC002 | *C. glutamicum* RES167 Δ*pldP* | RES167 Δ*pldP* | (1) |
| CEK01 | *C. glutamicum* RES167 *tatA::tatA-halotag* | RES167 *tatA::tatA-halotag* | This study |
| CEK02 | *C. glutamicum* RES167 Δ*pldP tatA::tatA-halotag* | RES167 Δ*pldP tatA::tatA-halotag* | This study |
| CAD05 | *C. glutamicum* RES167 *tatA::tatA-mCherry* | RES167 *tatA::tatA-mCherry* | This study |
| CAD06 | *C. glutamicum* RES167 Δ*pldP tatA::tatA-mCherry* | RES167 Δ*pldP tatA::tatA-mCherry* | This study |

| | | | |
|---|---|---|---|
| CDMB01 | *C. glutamicum* RES167 pDMB-RR-0955-H6 | RES167 with pCLTON1-$P_{sod}$-*cg0955-H6* | This study |
| CDMB02 | *C. glutamicum* RES167 pDMB-KK-0955-H6 | RES167 with pCLTON1-$P_{sod}$-*KK-cg0955-H6* | This study |
| CDMB03 | *C. glutamicum* RES167 pDMB-RR-PhoD-H6 | RES167 with pCLTON1-$P_{sod}$-*phoD-H6* | This study |
| CDMB04 | *C. glutamicum* RES167 pDMB-KK-PhoD-H6 | RES167 with pCLTON1-$P_{sod}$-*KK-phoD-H6* | This study |
| CDMB05 | *C. glutamicum* RES167 Δ*pldP* pDMB-RR-0955-H6 | RES167 Δ*pldP* with pCLTON1-$P_{sod}$-*cg0955-H6* | This study |
| CDMB06 | *C. glutamicum* RES167 Δ*pldP* pDMB-KK-0955-H6 | RES167 Δ*pldP* with pCLTON1-$P_{sod}$-*KK-cg0955-H6* | This study |
| CDMB07 | *C. glutamicum* RES167 Δ*pldP* pDMB-RR-PhoD-H6 | RES167 Δ*pldP* with pCLTON1-$P_{sod}$-*phoD-H6* | This study |
| CDMB08 | *C. glutamicum* RES167 Δ*pldP* pDMB-KK-PhoD-H6 | RES167 Δ*pldP* with pCLTON1-$P_{sod}$-*KK-phoD-H6* | This study |
| CEK03 | *C. glutamicum* RES167 pEKEx2-*tatSP-mNeonGreen* | RES167 with IPTG-inducible pEKEx2-*tatSP-mNeonGreen* | This study |
| CEK04 | *C. glutamicum* RES167 Δ*pldP* pEKEx2-*tatSP-mNeonGreen* | RES167 Δ*pldP* with IPTG-inducible pEKEx2-*tatSP-mNeonGreen* | This study |

9

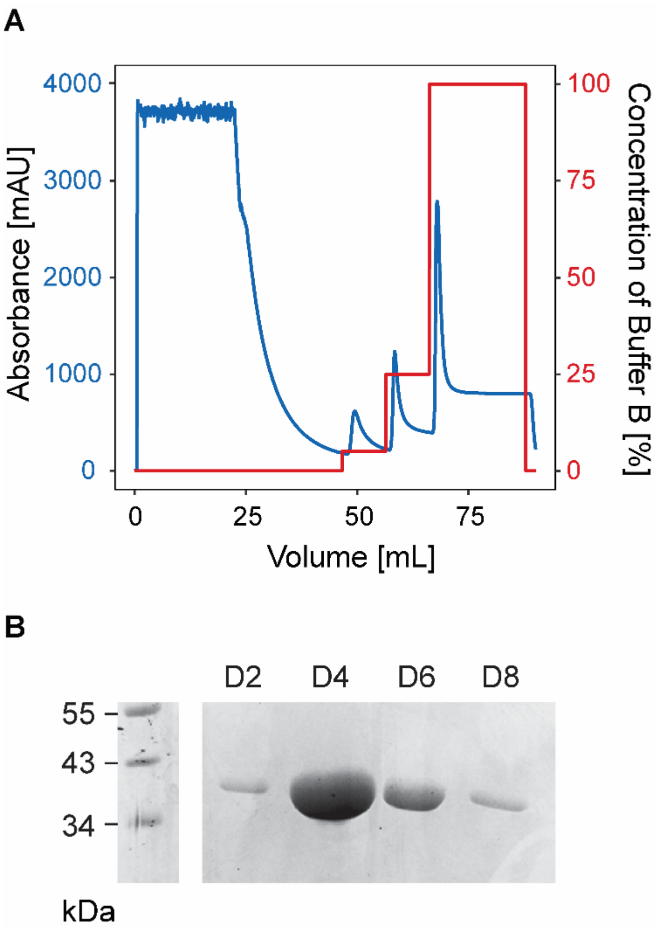


**Figure S1: Purification of PldP via Ni-NTA affinity chromatography. (A)** PldP is eluted in the presence of imidazole [500 mM] in Buffer B. Absorbance at 280 nm is depicted in blue colour, while the concentration of Buffer B – in red. **(B)** Eluted fractions D2, D4, D6 and D8 were collected and analyzed via SDS-PAGE. The fractions correspond to the third peak of the absorbance curve at 100% of Buffer B. The fractions D3, D4 and D5 were pulled together and used in further biochemical analysis. mAU – milli-absorbance units at 280 nm.

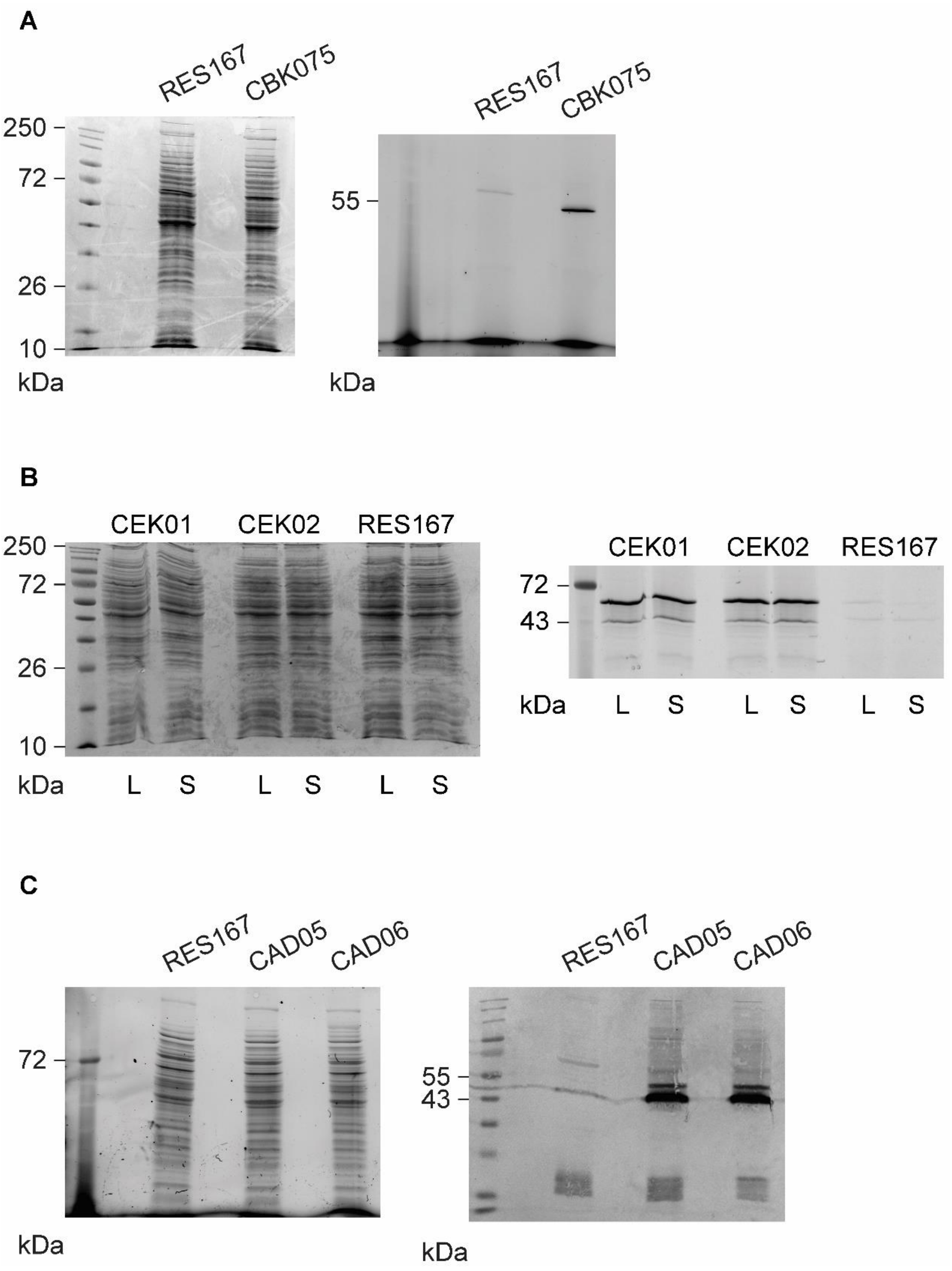


**Figure S2: Protein fusions are confirmed via in-gel fluorescence and Western blotting. (A)** PldP-eYFP in *C. glutamicum* RES167 *pldP::pldP-eYFP* (CBK075). Cells were grown in BHI medium, and their $OD_{600}$ were adjusted to 15 in 1 mL. Cells were resuspended in PBS buffer [1X] and disrupted by sonication. After cell lysis and centrifugation for 10 min at 14,100 rcf, 40 µL of the supernatant were mixed with SDS loading dye [1X] and incubated for 20 min at RT. Proteins were separated via SDS-PAGE [10 µL of the sample per well]. First image –

Coomassie brilliant blue-stained gel. Second image – in-gel fluorescence. Molecules of eYFP were excited using Blue Epi illumination, and the emission was captured using a 530/28 filter (eYFP excitation = 513 nm; emission = 527 nm). Size of proteins: eYFP – 27 kDa; PldP – ~31.57 kDa (predicted *in silico*). **(B)** TatA-HaloTag-TMR in *C. glutamicum* RES167 *tatA::tatA-halotag* (CEK01) and *C. glutamicum* RES167 Δ*pldP tatA::tatA-halotag* (CEK02). Cells were grown in BHI, and their $OD_{600}$ were adjusted to 15 in 1 mL. Cells were stained with the HaloTag ligand TMR [50 nM] and then washed in PBS [1X]. Cells were disrupted by sonication, and the samples were prepared as mentioned above. First image – Coomassie brilliant blue-stained gel. Second image – in-gel fluorescence. Molecules of TMR were excited using Green Epi illumination, and the emission was captured using a 605/50 filter (TMR excitation = 557 nm; emission = 576 nm). Size of proteins: HaloTag – 33 kDa; TatA – 12 kDa. L – cell lysate; S – supernatant. **(C)** TatA-mCherry in *C. glutamicum* RES167 *tatA::tatA-mCherry* (CAD05) and *C. glutamicum* RES167 Δ*pldP tatA::tatA-mCherry* (CAD06). Samples were prepared as described above. First image – gel with 2,2,2-trichloroethanol (TCE). Second image – polyvinylidene difluoride (PVDF) membrane after Western blotting, incubation with antibodies and development. Primary antibodies – anti-mCherry antibodies produced in the rabbit. Secondary antibodies – anti-rabbit alkaline phosphatase antibodies produced in the goat (Sigma-Aldrich). Size of proteins: mCherry – 28.8 kDa; TatA – 12 kDa. In every image, the first well is Color Prestained Protein Standard, Broad Range (10-250 kDa; NEB).

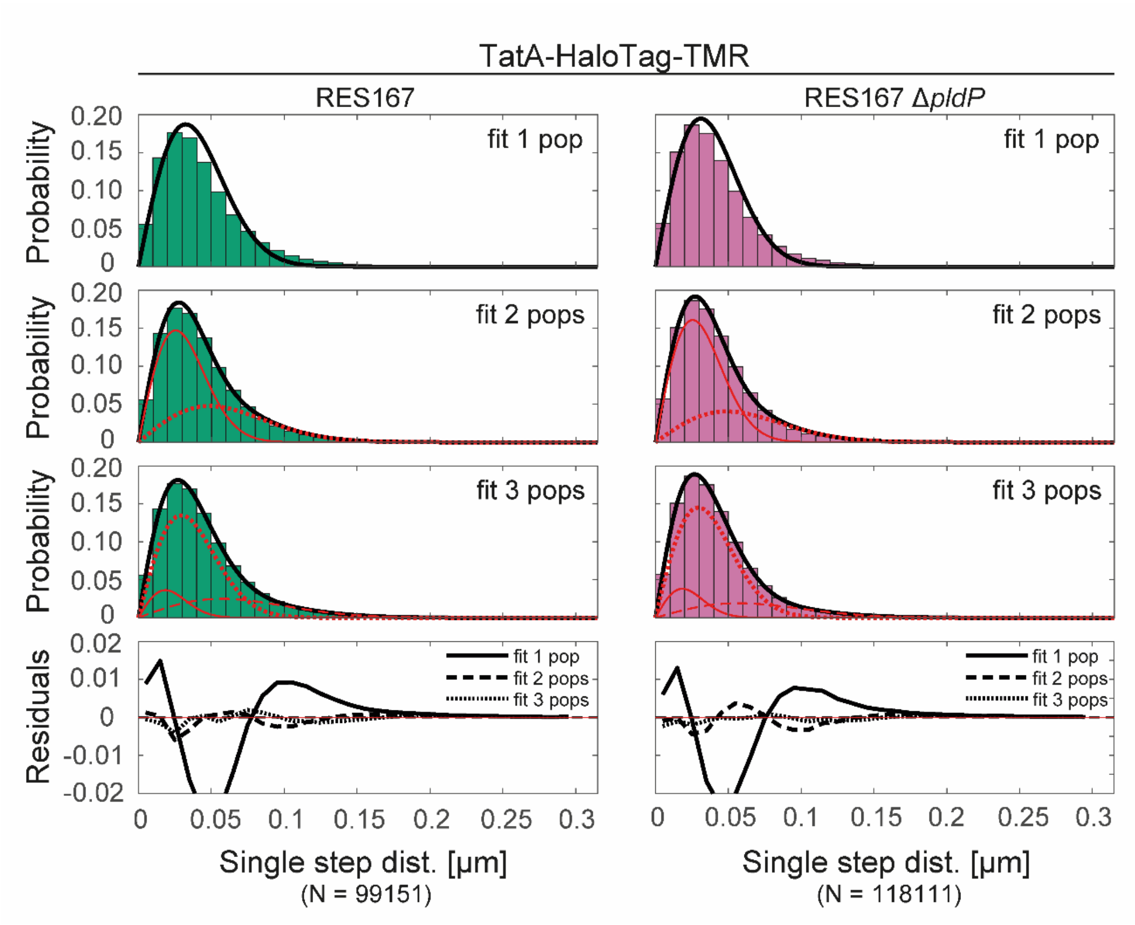


**Figure S3: Jump distance (JD) analysis of TatA-HaloTag-TMR.** JD analysis was performed for *C. glutamicum* RES167 *tatA::tatA-halotag* (CEK01) and *C. glutamicum* RES167 Δ*pldP tatA::tatA-halotag* (CEK02). JD analysis was based on the diffusion constants and population fractions of TatA-HaloTag-TMR, which were determined by fitting the cumulative distribution function (CDF) of the square displacements (SQD) (see Materials and Methods). Single populations are shown as red curves; the sum of the populations – as black curves. One-, two- and three-component models were fitted to the data. The residual plots show that there is no difference between two- and three-component model fitting; therefore, the two-component model is preferred. This results in two relative populations of TatA-HaloTag-TMR molecules – confined and slow mobile.

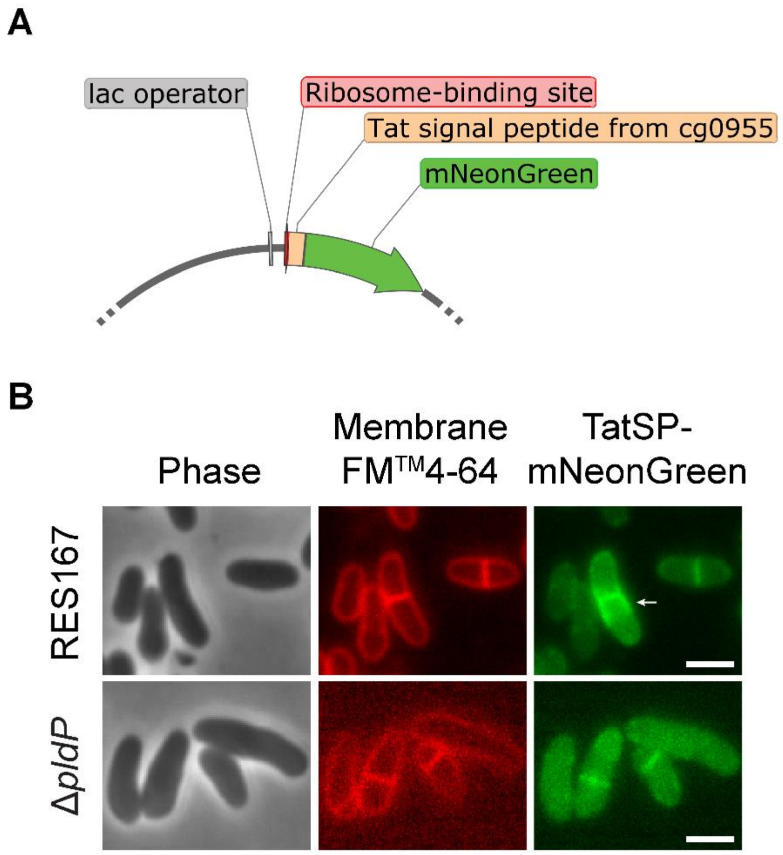


**Figure S4: Distribution of mNeonGreen fused to the Cg0955 Tat signal peptide is affected by *pldP* deletion. (A)** Fragment of the pEKEx2 plasmid containing *mNeonGreen* fused to the Tat signal peptide from *cg0955*. Two fragments were inserted into the multiple cloning site: Tat signal peptide sequence from *cg0955* (90 bp), together with an upstream ribosome-binding site (109 bp) and *mNeonGreen* (711 bp). The overexpression of *tatSP-mNeonGreen* is induced by IPTG [10 µM]. **(B)** TatSP-mNeonGreen is localized to the septum and neighbouring membrane regions in *C. glutamicum* RES167 pEKEx2-*tatSP-mNeonGreen* (CEK03). White arrow indicates the fluorescence at the lateral membrane regions close to the septum. In the case of *C. glutamicum* RES167 Δ*pldP* pEKEx2-*tatSP-mNeonGreen* (CEK04), the mNeonGreen fluorescence is either observed at the septum or dispersed throughout the cytoplasm. In both strains, the fluorescence of mNeonGreen at the septum is often observed after the membrane separation of daughter cells has already taken place. The cell populations are heterogeneous in relation to the mNeonGreen fluorescence. In particular, in CEK03, some cells show a higher fluorescence signal compared to others. The cell membranes were stained with FM$^{TM}$4-64 dye. Exposure time: FM$^{TM}$4-64 – 200 ms; mNeonGreen – 200 ms.
